# Widespread retention of ohnologs in key developmental gene families following whole genome duplication in arachnopulmonates

**DOI:** 10.1101/2020.07.10.177725

**Authors:** Amber Harper, Luis Baudouin Gonzalez, Anna Schönauer, Ralf Janssen, Michael Seiter, Michaela Holzem, Saad Arif, Alistair P. McGregor, Lauren Sumner-Rooney

## Abstract

Whole genome duplications have occurred multiple times during animal evolution, including in lineages leading to vertebrates, teleosts, horseshoe crabs and arachnopulmonates. These dramatic events initially produce a wealth of new genetic material, generally followed by extensive gene loss. It appears, however, that developmental genes such as homeobox genes, signalling pathway components and microRNAs are frequently retained as duplicates (so called ohnologs) following whole-genome duplication. These not only provide the best evidence for whole-genome duplication, but an opportunity to study its evolutionary consequences. Although these genes are well studied in the context of vertebrate whole-genome duplication, similar comparisons across the extant arachnopulmonate orders are patchy. We sequenced embryonic transcriptomes from two spider species and two amblypygid species and surveyed three important gene families, Hox, Wnt and frizzled, across these and twelve existing transcriptomic and genomic resources for chelicerates. We report extensive retention of putative ohnologs, including amblypygids, further supporting the ancestral arachnopulmonate whole-genome duplication. We also find evidence of consistent evolutionary trajectories in Hox and Wnt gene repertoires across three of the five arachnopulmonate orders, with inter-order variation in the retention of specific paralogs. We identify variation between major clades in spiders and are better able to reconstruct the chronology of gene duplications and losses in spiders, amblypygids, and scorpions. These insights shed light on the evolution of the developmental toolkit in arachnopulmonates, highlight the importance of the comparative approach within lineages, and provide substantial new transcriptomic data for future study.

## INTRODUCTION

The duplication of genetic material is an important contributor to the evolution of morphological and physiological innovations (Ohno 1970; Zhang 2003). The most dramatic example of this is whole genome duplication (WGD), when gene copy numbers are doubled and retained paralogs (ohnologs) can then share ancestral functions (subfunctionalization) and/or evolve new roles (neofunctionalization) (Ohno 1970; Force et al. 1999; Lynch and Conery 2000). The occurrence of two rounds (2R) of WGD in the early evolution of vertebrates has long been associated with their taxonomic and morphological diversity (e.g. Ohno 1970; Holland et al. 1994; Dehal and Boore 2005; Holland 2013), and a subsequent 3R in teleosts is frequently linked to their success as the most diverse vertebrate group (e.g. Meyer and Schartl 1999; Glasauer and Neuhauss 2014). However, this remains controversial and difficult to test (Donoghue and Purnell 2005) and in several animal lineages there is no clear association between WGD and diversification (Mark Welch et al. 2008; Flot et al. 2013; Havlak et al. 2014; Kenny et al. 2016; Nong et al. 2020). Along with vertebrates, chelicerates also appear to be hotspots of WGD, with up to three rounds reported in horseshoe crabs (Havlak et al. 2014; Kenny et al. 2016; Nong et al. 2020), one in the ancestor of arachnopulmonates (spiders, scorpions, and their allies) (Schwager et al. 2017), and potentially two further rounds within the spider clade Synspermiata (Král et al. 2019). Chelicerates demonstrate a highly variable body plan, occupy a wide range of habitats and ecological niches, and have evolved a variety of biologically important innovations such as venoms and silks (Schwager et al. 2015). They therefore offer an excellent opportunity for comparison with vertebrates concerning the implications of WGD for morphological and taxonomic diversity, and genome evolution in its wake.

The house spider *Parasteatoda tepidariorum* has emerged as a model species to study the impacts of WGD on arachnid evolution and development. Genomic and functional developmental studies have found retained ohnologs of many important genes, with evidence for neo- and subfunctionalization compared to single-copy orthologs in arachnids lacking WGD (Janssen et al. 2015; Leite et al. 2016; Turetzek et al. 2016; Schwager et al. 2017; Turetzek et al. 2017; Leite et al. 2018; Baudouin-Gonzalez et al. 2021). Work on the scorpions *Centruroides sculpturatus* and *Mesobuthus martensii* has consistently complemented findings in *P. tepidariorum*, with genomic studies recovering many ohnologs retained in common with spiders (Di et al. 2015; Sharma et al. 2015; Leite et al. 2018). In the past few years, several additional spider genomes have become available, providing an opportunity to get a more detailed view of genome evolution following WGD. Although synteny analysis remains the gold standard for the identification of ohnologs, the required chromosome-level genomic assemblies remain relatively scarce. Work on the *P. tepidariorum, C. sculpturatus* and *M. martensii* genomes has been complemented by targeted studies of individual gene families and transcriptomic surveys (Schwager et al. 2007; Sharma et al. 2012; Leite et al. 2018; Gainett and Sharma 2020). Combined with phylogenetic analyses, the identification of duplications can provide evidence of WGD events and their timing in arachnid evolution. Although transcriptomes can yield variant sequences of individual genes, from different alleles or individuals in mixed samples, these are generally straight-forward to filter out from truly duplicated loci owing to substantial sequence divergence in the latter. They also offer the double-edged sword of capturing gene expression, rather than presence in the genome; pseudogenized or silenced duplicates are not detected, but neither are functional genes if they are not expressed at the sampled timepoint or tissue. Such studies have produced strong additional evidence for an ancestral WGD, with patterns of duplication coinciding with our expectations for arachnopulmonate ohnologs (Clarke et al. 2014; Clarke et al. 2015; Sharma et al. 2015; Turetzek et al. 2017; Leite et al. 2018; Gainett and Sharma 2020; Gainett et al. 2020).

Comparison of WGD events among arachnopulmonates, horseshoe crabs and vertebrates indicates that despite extensive gene loss following duplication events, certain gene families are commonly retained following duplication (Holland et al. 1994; Schwager et al. 2007; Kuraku and Meyer 2009; Di et al. 2015; Sharma et al. 2015; Kenny et al. 2016; Leite et al. 2016; Schwager et al. 2017; Leite et al. 2018). These typically include genes from the conserved developmental ‘toolkit’ of transcription factors (TFs), cell signalling ligands and receptors, and microRNAs (Erwin 2009). Among these, several have stood out as focal points in the study of gene and genome duplications. The Hox group of homeobox genes regulate the identity of the body plan along the antero-posterior axis of all bilaterian animals (McGinnis and Krumlauf 1992; Abzhanov et al. 1999; Carroll et al. 2005; Pearson et al. 2005; Hueber and Lohmann 2008; Holland 2013). Four clusters of these key developmental genes were retained after 1R and 2R in vertebrates (Holland et al. 1994; Meyer and Schartl 1999; Kuraku and Meyer 2009; Pascual-Anaya et al. 2013), and the arachnopulmonate WGD is evident in the almost universal retention of Hox gene duplicates in sequenced genomes, with two ohnologs of all ten arthropod Hox genes in the scorpion *M. martensii* (Di et al. 2015; Leite et al. 2018), all except *Hox3* being represented by two copies in *C. sculpturatus* (Leite et al. 2018), and all except *fushi tarazu* (*ftz*) in *P. tepidariorum* (Schwager et al. 2017). Systematic studies of Hox gene expression patterns in the latter demonstrated that all nine pairs of Hox paralogs exhibit signs of sub- or neofunctionalization (Schwager et al. 2017). This high level of retention and expression divergence lends strong support to the importance of Hox gene duplication in the evolution of the arachnopulmonate body plan, and further consolidates the position of this family as a key indicator of WGD. Additionally, arachnids lacking WGD, such as ticks, mites, and harvestmen, exhibit single copies of the Hox genes, with no evidence for duplication via other routes (Grbić et al. 2011; Leite et al. 2018; Gainett et al. 2021).

In addition to TFs, the ligands and receptors of some signalling pathways of the developmental toolkit (e.g. Hedgehog, Wnt, TGF-ß, NHR) also demonstrate higher copy numbers in vertebrates and other groups subject to WGD, including arachnopulmonates (Holland et al. 1994; Meyer and Schartl 1999; Shimeld 1999; Pires-daSilva and Sommer 2003; Cho et al. 2010; Janssen et al. 2010; Hogvall et al. 2014; Janssen et al. 2015). The Wnt signalling pathway plays many important roles during animal development, including segmentation and patterning of the nervous system, eyes and gut (Erwin 2009; Murat et al. 2010). In the canonical pathway, Wnt ligands bind to transmembrane receptors, such as Frizzled, to trigger translocation of ß-catenin to the nucleus and mediate regulation of gene expression (Cadigan and Nusse 1997; Hamilton et al. 2001; Logan and Nusse 2004; van Amerongen and Nusse 2009). There are thirteen subfamilies of Wnt genes found in bilaterians, as well as multiple receptor families and downstream components. In contrast to the extensive retention of Hox ohnologs following WGD, Wnt duplicates in *P. tepidariorum* appear to be restricted to *Wnt7* and *Wnt11*, with the remaining eight subfamilies represented by single genes (Janssen et al. 2010). However, these are the only reported Wnt gene duplications in arthropods despite several recent surveys (Bolognesi et al. 2008; Murat et al. 2010; Hayden and Arthur 2013; Meng et al. 2013; Hogvall et al. 2014; Janssen and Posnien 2014; Holzem et al. 2019), and beyond *P. tepidariorum* no other arachnopulmonates have been systematically searched. Similarly, duplications within the four *frizzled* gene subfamilies appear to be restricted to arachnopulmonates among arthropods, wherein only *fz4* is duplicated in both *P. tepidariorum* and *M. martensii* (Janssen et al. 2015).

Several Wnt families have also been retained after the 1R and 2R events in vertebrates, for example there are two copies each of *Wnt2, Wnt3, Wnt5, Wnt7, Wnt8, Wnt9*, and *Wnt10* in humans (Miller 2001; Janssen et al. 2010). However, no subfamilies are represented by three or four copies in humans and so there is some consistency with arachnopulmonates in that the Wnts may be more conservative markers of WGD, to be used in combination with Hox and other homeobox genes.

The extensive and consistent retention of key developmental genes like Hox genes apparent in *P. tepidariorum* and *C. sculpturatus*, and Wnt genes in *P. tepidariorum*, strongly support the occurrence of an ancestral WGD in arachnopulmonates. However, data are only available for a handful of species so far, resulting in very patchy taxonomic sampling. For example, only *P. tepidariorum* and *Pholcus phalangioides* have been comprehensively surveyed for homeobox genes among spiders (Leite et al. 2018), omitting the large and derived retrolateral tibial apophysis (RTA) clade, which includes jumping spiders, crab spiders and other free hunters, and the systematic identification of Wnt genes has been restricted to only *P. tepidariorum*. Spiders and scorpions are by far the most speciose of the arachnopulmonates, and there may be additional diversity in their repertoires of these important developmental gene families of which we are not yet aware.

In addition, and perhaps more urgently, only two of the five arachnopulmonate lineages have dominated the field thus far; sufficient genomic information for comparison is lacking beyond spiders and scorpions. Also represented in Arachnopulmonata are the amblypygids (whip spiders), relatively understudied and enigmatic animals comprising around 190 extant species. They exhibit highly derived morphology of the pedipalps, which are adapted to form raptorial appendages, and of the first pair of walking legs, which are antenniform and can comprise more than 100 segments (Weygoldt 2009). Despite the scarcity of transcriptomic or genomic data for amblypygids (whip spiders) (Gainett and Sharma 2020; Gainett et al. 2020 for recent advances), their widely accepted position within Arachnopulmonata implies that they were also subject to an ancestral WGD. A recent survey of the *Phrynus marginemaculatus* transcriptome supported this in the recovery of multiple duplicate Hox and leg gap genes (Gainett and Sharma 2020). Particularly given the derived nature of their appendages, this group could shed substantial light on genomic and morphological evolution following WGD.

To better understand the genomic consequences of WGD in a greater diversity of arachnopulmonate lineages, we sequenced *de novo* embryonic transcriptomes from two spiders belonging to the derived RTA clade and two amblypygids. We surveyed Hox, Wnt and frizzled genes in these species and existing genomic and transcriptomic resources for comparison with other arachnids, both with and without an ancestral WGD, improving sampling at both the order and sub-order levels.

## MATERIALS AND METHODS

### Embryo collection, fixation and staging

Embryos of mixed ages were collected from captive females of the amblypygids *Charinus acosta* (Charinidae; parthenogenetic, collected at one day, one month and two months after the appearance of egg sacs; equivalent to approximately 1%, 30% and 60% of development) and *Euphrynichus bacillifer* (Neoamblypygi: Phrynichidae; mated, collected at approximately 30% of development), the wolf spider *Pardosa amentata* (collected in Oxford, UK, equivalent to stages 11/12 in *P. tepidariorum*) and mixed stage embryos of the jumping spider *Marpissa muscosa* (kindly provided by Philip Steinhoff and Gabriele Uhl) and stored in RNAlater. *Phalangium opilio* were collected in Uppsala, Sweden, and developmental series of embryos ranging from egg deposition to the end of embryogenesis were collected for sequencing.

### Transcriptomics

Total RNA was extracted from mixed aged embryos, pooled by species, of *Charinus acosta, Euphrynichus bacillifer, Pardosa amentata* and *Marpissa muscosa* using QIAzol lysis reagent according to the manufacturer’s instructions (Qiagen). Illumina libraries were prepared using a TruSeq RNA kit (including polyA selection) and sequenced on the NovaSeq platform (100bp PE, Edinburgh Genomics). The quality of raw reads was assessed using FastQC v0.11.9 (Andrews 2010), erroneous k-mers were corrected using rCorrector (default settings; Song and Florea 2015), and unfixable read pairs (from low-expression homolog pairs or containing too many errors) were discarded using a custom Python script (available at https://github.com/harvardinformatics/TranscriptomeAssemblyTools/blob/master/FilterUncorrectabledPEfastq.py courtesy of Adam Freeman). Adapter sequences were identified and removed and low quality ends (phred score cut-off = 5) trimmed using TrimGalore! v0.6.5 (available at https://github.com/FelixKrueger/TrimGalore). *De novo* transcriptome assembly was performed using only properly paired reads with Trinity v2.10.0 (Haas et al. 2013) using default settings. Transcriptome completeness was evaluated on the longest isoform per gene using BUSCO v4.0.2 (77) along with the arachnid database (arachnida_odb10 created on 2019-11-20; 10 species, 2934 BUSCOs) and the arthropod database (arthropoda_odb10 created on 2019-11-20; 90 species, 1013 BUSCOs). Reads are available on SRA (BioProject PRJNA707377) and assembled transcriptomes are available at (XXXXX).

### Identification of gene candidates

To identify Hox, Wnt and frizzled gene candidates across chelicerates, we performed BLAST searches (p-value 0.05) against existing genomic and transcriptomic resources and the four new embryonic transcriptomes generated in this study. Hox, Wnt and Frizzled peptide sequences previously identified in *P. tepidariorum* and *Ixodes scapularis* were reciprocally blasted against the respective NCBI proteomes and the top hit was selected (Supplementary File 1) (Janssen et al. 2010; Janssen et al. 2015; Schwager et al. 2017; Leite et al. 2018). Hox protein sequences previously identified in *C. sculpturatus* (Schwager et al. 2017) and *P. opilio* (Leite et al. 2018) were reciprocally blasted against the respective NCBI proteomes and the top hit was selected (Supplementary File 1). These sequences along with the Hox and Fz peptide sequences identified in *M. martensii* genome were used as query sequences in our analysis (Di et al. 2015; Janssen et al. 2015).

BLASTP searches were performed against the NCBI proteomes of *Stegodyphus dumicola*, *C. sculpturatus*, *Tetranychus urticae* and *Limulus polyphemus*. TBLASTN searches were performed against *C. acosta*, *E. bacillifer*, *P. amentata* and *M. muscosa* (this study); the transcriptomes of *P. phalangioides* (Turetzek, Torres-Oliva, Kaufholz, Prpic, Posnien, in preparation), *Phoxichilidium femoratum* (Ballesteros et al. 2021) and *P. opilio* (PRJNA236471); and the genomes of *Pardosa pseudoannulata* (PRJNA512163), *Acanthoscurria geniculata* (PRJNA222716), *M. martensii* (CDS sequence file; Cao et al. 2013). We then predicted the peptide sequences using the Translate ExPASy online tool (https://web.expasy.org/translate/<u>; default settings</u>). Protein sequence identity was confirmed by reciprocal BLAST against the NCBI database and the construction of maximum likelihood trees. Where more than one sequence was identified as a potential candidate for a single gene, nucleotide and protein alignments were inspected to eliminate the possibility that they were isoforms, individual variants, or fragments of the same gene. Only the longest isoforms and gene fragments were selected for phylogenetic analysis (Supplementary File 1). All Hox sequences identified contain a complete or partial homeodomain (Supplementary Files 2-3). The TBLASTN search in *P, phalangioides* identified a *Ubx* gene not reported previously by Leite et al. (2018). The Trinity transcript accession numbers of all identified sequences, NCBI protein accession and other sequence identifiers used in the subsequent phylogenetic analysis are found in Supplementary File 1.

Due to high levels of fragmentation in *P. opilio*, multiple non-overlapping fragments were found to align with query Wnt sequences. To verify the identity and relationship of these fragments, primers were designed against the 5’-most and 3’-most ends of the aligned series (see Supplementary File 1). Total RNA was extracted using Trizol (Invitrogen) according to the manufacturer’s instructions and cDNA produced using the SuperScriptII first-strand synthesis system (Invitrogen) for RT-PCR using the designed primer pairs. PCR products were sequenced by Macrogen Europe to confirm predicted combinations of fragmented transcripts (see Supplementary File 1).

### Phylogenetic analysis

Hox, Wnt and Frizzled protein predictions were retrieved from NCBI for the insects *Drosophila melanogaster*, *Bombyx mori*, *Tribolium castaneum*; the crustacean *Daphnia pulex*; and the onychophoran *Euperipatoides kanangrensis* (Supplementary File 1). The Hox, Wnt and Frizzled peptide sequences for the myriapod *Strigamia maritima* were retrieved from Chipman et al. (2014) (Supplementary File 1). Alignments were performed in Clustal Omega using default parameters (Larkin et al. 2002; Sievers et al. 2011), with the exception of the full Hox protein sequences, which were aligned using COBALT (Papadopoulos and Agarwala 2007). Maximum likelihood trees were generated from whole-sequence alignments to assign sequences to families and study the relationship between candidate duplicates. Phylogenetic analyses were performed in IQ-Tree (v2.0.3, Nguyen et al. 2015) using ModelFinder to identify optimal substitution models (JTT+F+R9 for full Hox sequences, LG+G4 for Hox homeodomain sequences, GTR+F+I+G4 for Hox homeobox sequences, LG+R8 for Wnt, JTT+R8 for Fz; Kalyaanamoorthy et al. 2017) and 100,000 bootstrap replicates. Trees were visualized in FigTree v.1.4.4 (http://tree.bio.ed.ac.uk/software/figtree/) and tidied in Adobe Illustrator. Hox sequences were additionally analysed using RaXML v8 (Stamatakis 2014), using the same substitution models chosen by Iqtree and the automatic bootstopping algorithm (Pattengale et al. 2009). Alignments are provided in Supplementary Files 2-4 and 9-11.

## RESULTS

### Transcriptome assemblies

To further study the outcomes of WGD in the ancestor of arachnopulmonates we carried out RNA-Seq on embryos of two further spider species, *P. amentata* and *M. muscosa*, and two species of amblypygids, *C. acosta* and *E. bacillifer*.

RNA-Seq for the four species produced between 222,479,664 and 272,844,971 raw reads, reduced to 211,848,357 and 260,853,757 after processing. Trinity assembled between 184,142 and 316,021 transcripts in up to 542,344 isoforms (Table 1). Contig N50 ranged from 592 bp in *M. muscosa* to 978 bp in *E. bacillifer*, and from 1461 bp (*M. muscosa*) to 2671 bp (*E. bacillifer*) in the most highly expressed genes (representing 90% of total normalized expression) (Table 1).

**Table 1.**
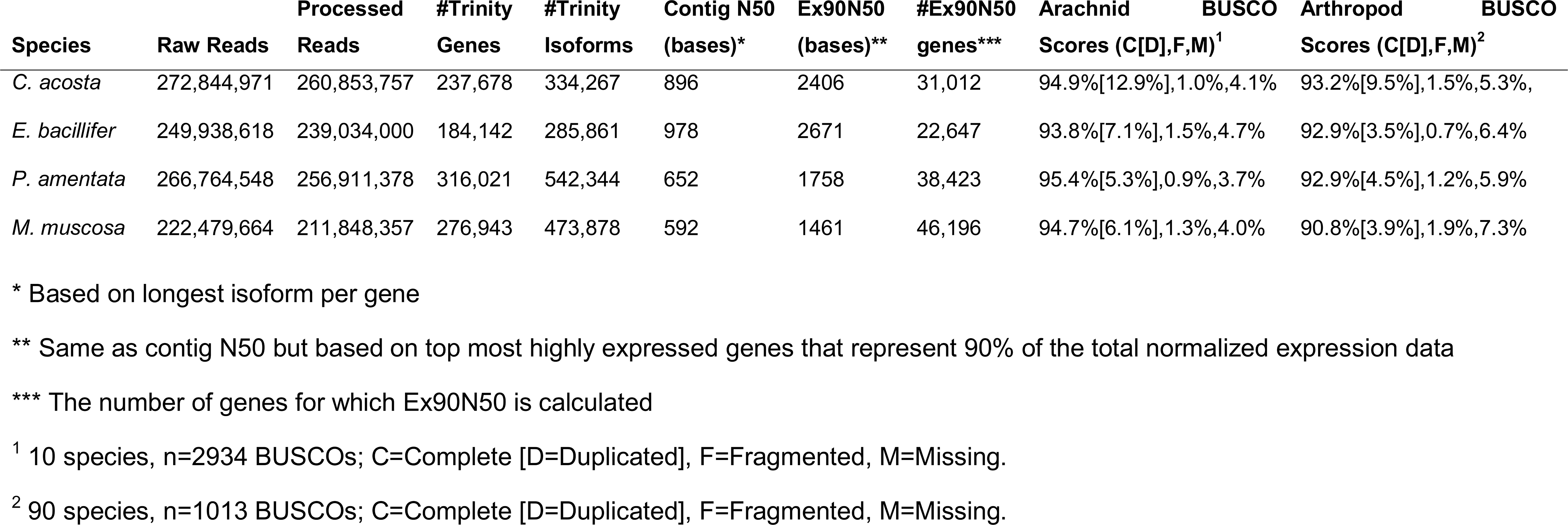
Assembly metrics for transcriptomes of Charinus acosta, Euphrynichus bacillifer, Marpissa muscosa and Pardosa amentata.

Transcriptomes were found to be between 83.7% (*C. acosta*) and 89.4% (*E. bacillifer*) complete according to BUSCO scores compared to the arthropod database, with between 3.5% and 9.5% duplicated BUSCOs. Compared to the arachnid databases, transcriptomes were 82%-90.1% complete for single-copy BUSCOs and contained between 5.3%-12.9% duplicated BUSCOs (Table 1).

To explore the extent of duplication in these arachnopulmonates we then surveyed the copy number of Hox, Wnt and Frizzled genes in their transcriptomes in comparison to other arachnids. It is important to note that the absence of genes recovered from transcriptomes does not eliminate the possibility that they are present in the genome, as the transcriptomes will only capture genes expressed at the relevant point in development. Mixed-stage embryonic samples (all except *E. bacillifer*) may yield more transcripts for the same reason.

### Hox repertoires and their origins

Spider Hox gene repertoires are largely consistent with *Parasteatoda tepidariorum*, which has two copies of all except *ftz* (Figure 1). There are several exceptions: single copies of *Hox3* in *Marpissa muscosa*, *Pardosa amentata* and *Pardosa pseudoannulata*, of *Sex combs reduced (Scr)* in *Acanthoscurria geniculata* and *Stegodyphus dumicola*, of *proboscipedia* (*pb*) in *A. geniculata*, and of *labial* (*lab*) in *Pholcus phalangioides* (Figure 1). Although we found two *AbdB* candidates in *Pardosa pseudoannulata,* one of these did not contain a homeodomain and was excluded from phylogenetic analyses. In contrast to Leite et al. (2018) we recovered two copies of *Ubx* from *P. phalangioides*. The inclusion of additional sequences in this analysis also helped resolve issues in gene homologies previously presented in *P. phalangioides* by Leite et al. (2018).

**Figure 1.**
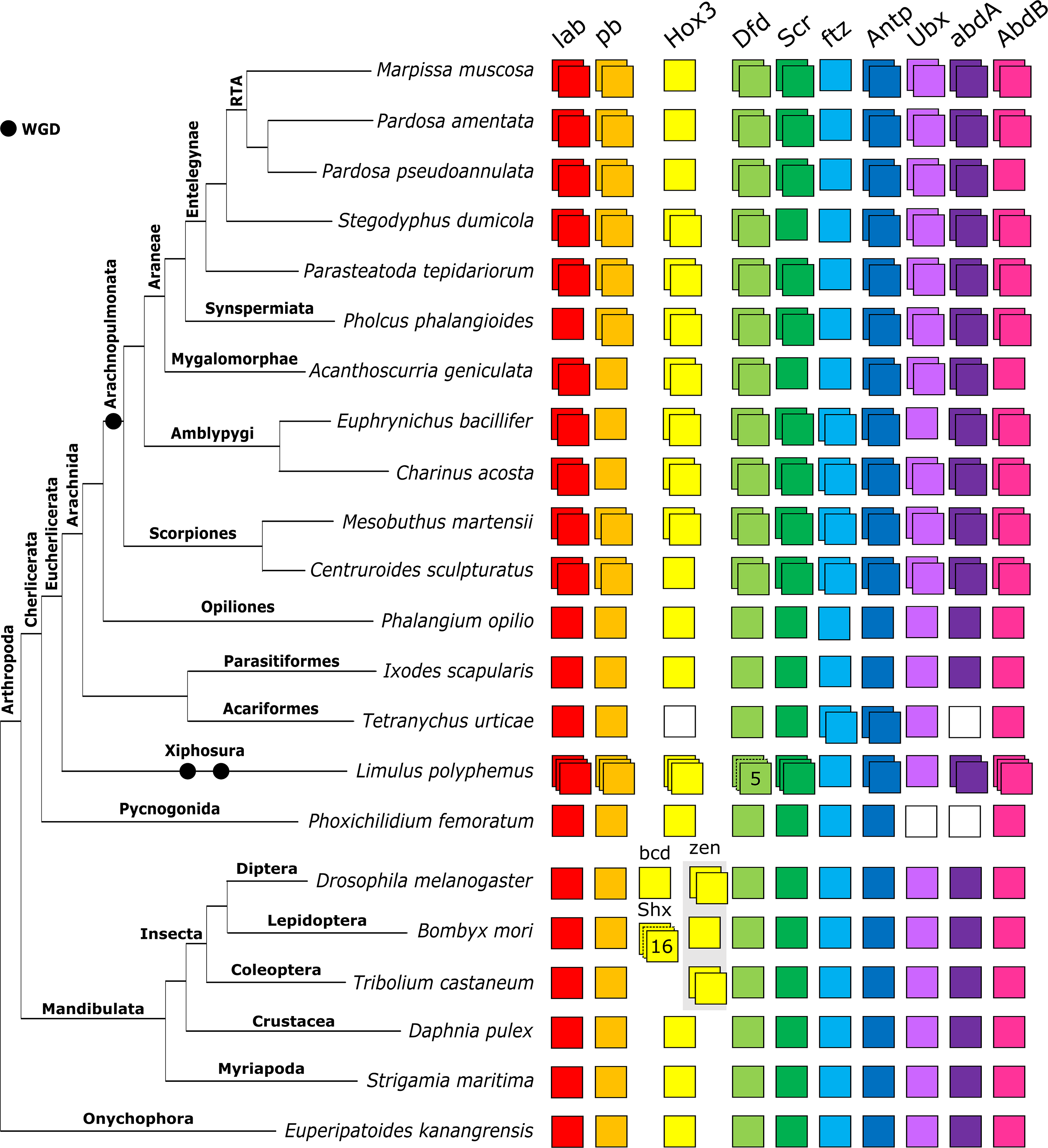
Repertoires of Hox genes in arachnids and other selected arthropods. Hox genes are represented by coloured boxes with duplicated Hox genes indicated by overlapping boxes. Figure includes Hox repertoires previously surveyed in the arachnids *P. tepidariorum, Centruroides sculpturatus*, *Mesobuthus martensii*, *Phalangium opilio*, *I. scapularis* (all genomes) and *Pholcus phalangioides* (embryonic transcriptome), the myriapod *Strigamia maritima* and the insects *Drosophila melanogaster, Tribolium castaneum* and *Bombyx mori*. The insect *Hox3* homolog *zen* has undergone independent tandem duplications in *T. castaneum* to yield *zen* and *zen2*; in cyclorrhaphan flies to yield *zen* and *bicoid*; and in the genus *Drosophila* to yield *zen2*. *Bombyx mori* is not representative of all species of ditrysian Lepidoptera, which typically possess four distinct *Hox3* genes termed Special homeobox genes (*ShxA*, *ShxB*, *ShxC* and *ShxD*) and the canonical *zen* gene.

Both amblypygids exhibit extensive duplication of Hox genes, with two copies recovered for all except for *pb* in *C. acosta* and *pb* and *Ubx* in *E. bacillifer* (Figure 1).

The scorpions *Mesobuthus martensii* and *Centruroides sculpturatus* exhibited similar Hox repertoires, with two copies recovered for all ten genes, except for a single copy of *Hox3* in *C. sculpturatus*.

Among the non-arachnopulmonate arachnids, *I. scapularis* and *P*. *opilio* exhibited no duplication of any Hox genes, in line with previous genomic surveys (Leite et al. 2018; Gainett et al. 2021). In the mite *Tetranychus urticae*, we did not identify *Hox3* and *abdA* candidates, consistent with Grbić et al. (2011) and Ontano et al. (2020). However, our BLAST search did not recover the *Antp*-2 copy previously identified by Grbić et al. (2011). The resolution of several sequences from *T. urticae* was variable; for example, previously identified *Tu-ftz-1*, *Tu-ftz-2*, *Tu-AbdB* and *Tu-Ubx* sequences (Grbić et al. 2011) did not resolve correctly in the full Hox tree (Figure 2) but did in the homeodomain and homeobox trees (Supplementary Files 5-8).

**Figure 2.**
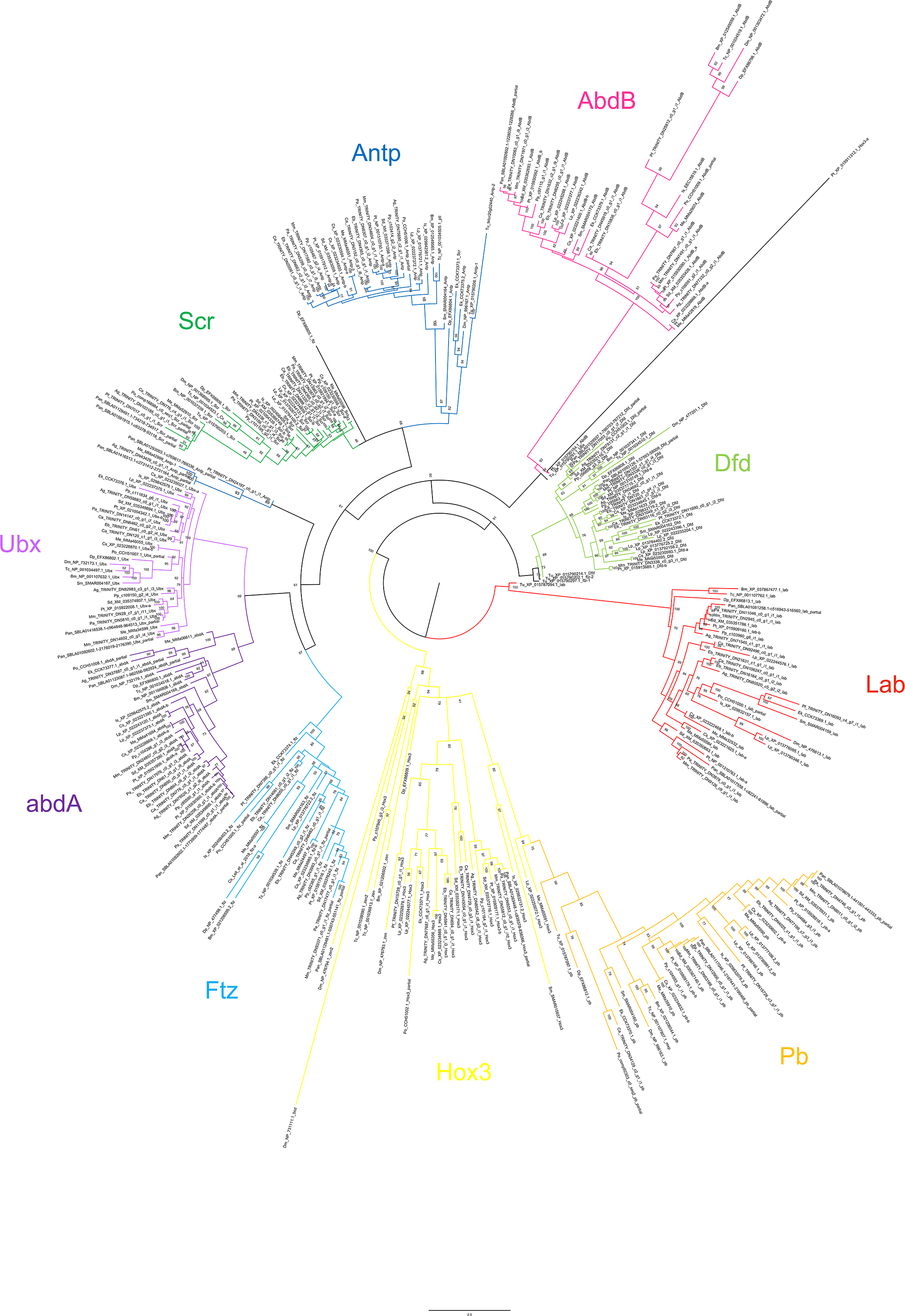
Maximum likelihood phylogeny of Hox amino acid sequences. The Hox genes are shown as different colours (after Figure 1). Panarthropods included: *Acanthoscurria geniculata* (Ag), *Bombyx mori* (Bm), *Centruroides sculpturatus* (Cs), *Charinus acosta* (Ca), *Drosophila melanogaster* (Dm), *Euperipatoides kanangrensis* (Ek), *Euphrynichus bacillifer* (Eb), *Ixodes scapularis* (Is), *Limulus polyphemus* (Lp), *Marpissa muscosa* (Mm), *Mesobuthus martensii* (Me), *Parasteatoda tepidariorum* (Pt), *Pardosa amentata* (Pa), *Pardosa pseudoannulata* (Pan), *Phalangium opilio* (Po), *Pholcus phalangioides* (Pp), *Phoxichildium femoratum* (Pf), *Stegodyphus dumicola* (Sd), *Strigamia maritima* (Sm), *Tetranychus urticae* (Tu), and *Tribolium castaneum* (Tc). Node labels indicate ultrafast bootstrap support values. See Supplementary File 1 for accession numbers, Supplementary File 2 for full amino acid sequence alignments.

We also surveyed two non-arachnid chelicerates, a pycnogonid *Phoxichilidium femoratum* and the horseshoe crab *Limulus polyphemus*, the latter of which has undergone multiple rounds of WGD independent of the arachnopulmonates (Kenny et al. 2016; Nong et al. 2020). We recovered single copies of all Hox genes except *Ubx* and *abdA*, which were not found, in *Phoxichilidium femoratum*, consistent with Ballesteros et al. (2021; Figure 1). *Limulus polyphemus* returned multiple copies of all Hox genes except *ftz* and *Ubx*, including three copies of *lab*, *pb*, *Hox3*, *Scr* and *AbdB,* five potential copies of *Dfd*, and two copies of *Antp* and *abdA* (Figure 1).

Full protein sequences produced the best supported phylogeny: six nodes returned support <50%. Two of these concerned the placement of outgroup (non-chelicerate) sequences and one concerned the deeper relationship of the Hox3 and Pb clades, which is beyond the scope of this study. The remaining three reflect uncertainty in the within-clade placement of chelicerate sequences (Figure 2). An additional ten nodes returned support of 50-60%, which we consider too weak to justify interpretation. Analyses using protein sequences from homeodomains only contained insufficient phylogenetic information to resolve within-clade relationships, but confirmed gene identity (Supplementary Files 5-6). Nucleotide sequences of homeodomains provided more within-clade resolution, but with lower support than full protein sequences (Supplementary Files 7-8). Using full protein sequences, we recover Antp as a paraphyletic group, albeit with low support (61%). The homeodomain phylogenies (both protein and nucleotide) confirm the identity of these sequences, indicating substantial sequence divergence outside of this conserved region in several species. The following discussion is based on the phylogeny of the full protein sequences (Figure 2).

Overall, the resolution of duplicate Hox sequences indicates that the majority are likely to be ohnologs. In many cases, duplicates form well-supported clades of orthologs, containing sequences from multiple Orders, suggesting a shared origin for duplication. The majority of Hox genes are present in duplicate across the arachnopulmonates, strengthening this pattern. Few duplicates form paralogous pairs within species, or paired clades of paralogs within lineages, as would be expected from lineage-specific duplications (although there are exceptions, such as scorpion Lab and Antp, Figure 2).

For both abdA and AbdB, sequences broadly formed two overall clades; although these were not well supported for AbdB (49-51%), relationships within them are still informative. In both cases, spider sequences formed two distinct and well-supported (>97%) clades that reflect overall phylogenetic relationships in their topology. The amblypygid sequences resolved as two ortholog pairs, one in each of the two overall clades. These are therefore strong candidates for ohnologs. Scorpion AbdB and abdA were not consistent with this pattern; in both cases, one pair of orthologs resolved together and one pair separately (Figure 2). Although this reflects a low likelihood of a lineage-specific duplication of abdA and AbdB in scorpions, it does not further clarify possible ohnolog relationships. Spider and scorpion Pb duplicates are candidate ohnologs, resolving in two well-supported (>99%) clades of orthologs whose topologies reflect phylogenetic relationships. Ftz duplicates in scorpions and amblypygids also resolved with orthologs of other arachnopulmonates, but one set of scorpion paralogs was placed with substantial uncertainty (support <60%). Most spider and amblypygid Antp sequences follow the pattern expected of ohnologs, forming two clades of orthologs, but three of the scorpion sequences formed a well-supported clade (91%) while the fourth resolved within the small group of sequences that fall outside the main Antp clade.

The origin of *Ubx*, *Dfd*, *Lab*, *Hox3* and *Scr* duplicates in arachnopulmonates is not clear from our phylogenetic analysis alone, with neither orthologs nor paralogs resolving together consistently, and topology that is a poor reflection of phylogeny. However, spider sequences broadly formed two clades with other arachnopulmonates, and synteny analysis in both *P. tepidariorum* (Schwager et al. 2017) and *Trichonephila antipodiana* (Fan et al. 2021) demonstrate clearly that Hox gene duplications therein are the result of WGD. The placement of the amblypygid and scorpion duplicates was more variable. In some cases these also appear to resolve as would be expected of ohnologs (e.g. amblypygid Hox3), but in others their placement could indicate lineage-specific duplication, although usually with low support (e.g. amblypygid Ubx).

It seems likely that having a single copy of *ftz* is common across all spiders (Figures 1-2), consistent with the loss of a duplicate in their common ancestor. A recently published high-quality genome from *Trichonephila antipodiana* also reported a single copy of *ftz* (Fan et al. 2021), as did another mygalomorph transcriptome, *Aphonopelma hentzi* (Ontano et al. 2020). The absence of *Hox3* duplicates in *M. muscosa, P. pseudoannulata* and *P. amentata* may indicate a lineage-specific loss in the RTA clade, which unites salticids, lycosids and their allies. Indeed, only one copy of *Hox3* was previously recovered in *Cupiennius salei*, which also belongs to the RTA clade (Schwager et al. 2007). Although the absence of a second copy in the embryonic transcriptomes could be attributed to failure to capture expression, both *C. salei* and *P. pseudoannulata* yielded single copies from genomic DNA, strengthening the case for a genuine loss. The *P. tepidariorum Hox3-A* sequence is highly divergent, and its expression is very weak (Schwager et al. 2017); this could indicate ongoing pseudogenisation. Other apparent losses of Hox gene duplicates are restricted to transcriptomes of individual species, with the exception of *Scr* in *Stegodyphus dumicola*, and could reflect failure to capture additional sequences.

The absence of a second copy of *pb* in both *Charinus acosta* and *Euphrynichus bacillifer*, which are distantly related within Amblypygi, suggests a loss in the common ancestor of amblypygids. Gainett and Sharma (2020) also recovered a single copy of *pb* in *Phrynus marginemaculatus*, but two copies in *Charinus israelensis*. However, one of these had an incomplete homeodomain (32 aa) that was identical to the other C-isr protein sequence; we are hesitant about its status as a duplicate. Embryos of *C. acosta* were collected at multiple stages of development, supporting the hypothesis that this is a genuine loss, rather than absence of expression at a particular developmental stage. However, the apparent absence of a *Ubx* duplicate in *E. bacillifer* could equivocally indicate lineage-specific loss or absence of expression at a single timepoint. In *P. tepidariorum* the two *Ubx* ohnologs are expressed most strongly in close succession, at stages 8.1 and 8.2 (Schwager et al. 2017), but this may not reflect relative expression patterns in amblypygids.

### Wnt repertoires and their origins

Consistent with *Parasteatoda tepidariorum* (Janssen et al. 2010), we found representatives of ten *Wnt* subfamilies in all surveyed spiders. The absence of *Wnt9* and *Wnt10* indicates their likely absence in the spider ancestor, while the absence of *Wnt3* is consistent with all other protostomes (Janssen et al. 2010; Murat et al. 2010; Hogvall et al. 2014). Two copies of *Wnt7* were retrieved from *Marpissa muscosa, Pardosa amentata, Acanthoscurria geniculata, Pholcus phalangioides, P. tepidariorum, Pardosa pseudoannulata and Stegodyphus dumicola*. All spiders except *A. geniculata* also yielded two copies of *Wnt11*. A second copy of *Wnt4,* which is absent in *P. tepidariorum,* was recovered from *P. amentata, M. muscosa, A. geniculata, S. dumicola* and *P. pseudoannulata*. We also found duplicates of *Wnt1/wg* in both *A. geniculata* and *P. pseudoannulata*, and a duplicate of *WntA* in *P. pseudoannulata*.

Representation of the Wnt subfamilies in the amblypygids is higher than any other chelicerate studied to date, including those with high-quality genome assemblies (Janssen et al. 2010; Hogvall et al. 2014; Holzem et al. 2019). We recovered transcripts from twelve out of thirteen subfamilies (missing *Wnt3*) and duplicates of *Wnt1/wg*, *Wnt4*, and *Wnt7* in both species (Figure 3). Additional duplicates of *Wnt6* and *Wnt11* were recovered from *Charinus acosta* (Figure 3).

**Figure 3.**
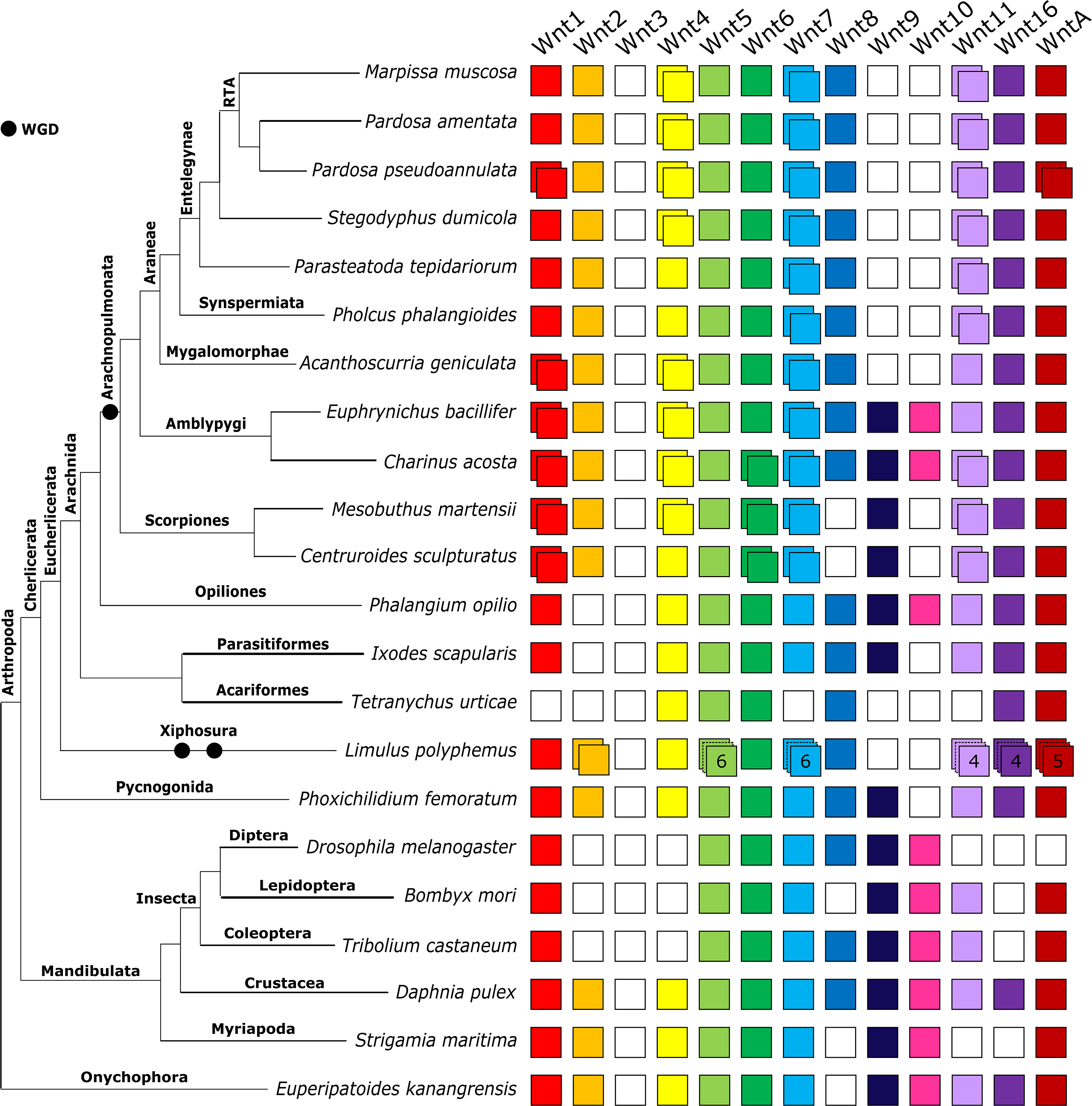
Repertoires of Wnt subfamilies in arthropods and an onychophoran. The Wnt subfamilies (1-11, 16 and A) are represented by coloured boxes with duplicated genes represented by overlapping boxes and putatively lost subfamilies indicated by white boxes. Figure includes Wnt repertoires recovered in this study and previously surveyed in the arachnids *Parasteatoda tepidariorum* and *Ixodes scapularis*; the insects *Drosophila melanogaster, Tribolium castaneum* and *Bombyx mori*; the crustacean *Daphnia pulex*; the myriapod *Strigamia maritima*; and the onychophoran *Euperpatoides kanangrensis*.

Representatives of ten Wnt subfamilies were found in the two scorpion genomes, with both missing *Wnt3*, *Wnt8* and *Wnt9*. Two copies of *Wnt1/wg*, *Wnt6*, *Wnt7* and *Wnt11* were retrieved in both species, with an additional copy of *Wnt4* in *M. martensii*.

We found no evidence of Wnt gene duplication in the non-arachnopulmonate arachnids (Figure 3). In the harvestman *Phalangium opilio* we recovered single copies of all except *Wnt2* and *Wnt3*. *Ixodes scapularis* was also missing *Wnt10* but was otherwise similar, whereas we also did not recover *Wnt1/wg*, *Wnt7*, *Wnt9*, or *Wnt11* in the mite *Tetranychus urticae*. In the non-arachnid chelicerates, *Phoxichilidium femoratum* and *Limulus polyphemus*, we found similar representation of the subfamilies, with all except *Wnt3* and *Wnt10* in *P. femoratum* and all except these and *Wnt9* in *L. polyphemus*. No duplicates were recovered in *P. femoratum*, but large numbers of duplicates were found in *L. polyphemus*. These included six potential copies of *Wnt5* and *Wnt7*, four of *Wnt11* and *Wnt16*, and five of *WntA*.

Only two nodes returned support <50%: one uniting Wnt8, Wnt9, Wnt10 and Wnt16 (39%), and one uniting Wnt1/wg, Wnt4, Wnt6 and Wnt11 (36%). These are positioned deep within the tree and concern the interrelationships of the Wnt subfamilies, which are beyond the scope of this study.

Most of the duplicate Wnt genes appear to be likely ohnologs; confirmation requires synteny analysis, but the relationships between paralogs resolved by phylogenetic analysis generally support duplications originating in the arachnopulmonate ancestor.

Duplicates of *Wnt7* were previously identified in *P. tepidariorum* (Janssen et al. 2010) and are recovered from all surveyed arachnopulmonates. The spider Wnt7 duplicates formed two clades (bootstrap ≥ 61%) suggesting the retention of ohnologs (Figure 4). The amblypygid Wnt7 ortholog pairs resolve as sisters to the spider clades, but support for these placements is lower (50-60%), and they display slightly higher sequence similarity between paralogs (73-74%) than the spiders (60-72%). The scorpion Wnt7 ortholog pairs formed their own separate clade with strong support (92%; Figure 4), possibly indicating a lineage-specific duplication. Nonetheless, the sequence divergence between the scorpion paralog pairs (68-70%) is similar to that between putative ohnologs in spiders (60-72%).

**Figure 4.**
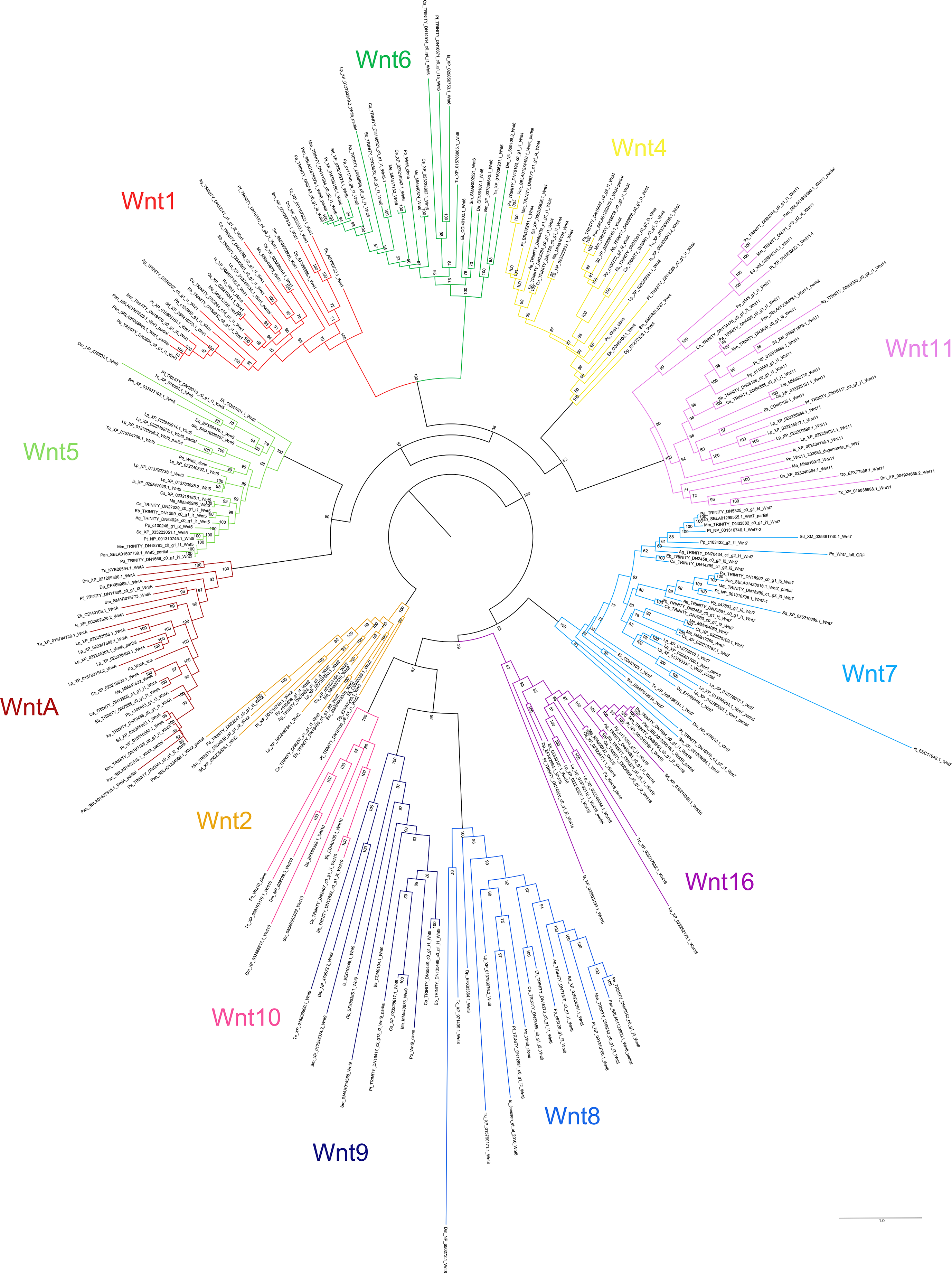
Maximum likelihood phylogeny of Wnt amino acid sequences. The 12 Wnt subfamilies are shown as different colours (after Figure 3). Panarthropods included: *Acanthoscurria geniculata* (Ag), *Bombyx mori* (Bm), *Centruroides sculpturatus* (Cs), *Charinus acosta* (Ca), *Drosophila melanogaster* (Dm), *Euperipatoides kanangrensis* (Ek), *Euphrynichus bacillifer* (Eb), *Ixodes scapularis* (Is), *Limulus polyphemus* (Lp), *Marpissa muscosa* (Mm), *Mesobuthus martensii* (Me), *Parasteatoda tepidariorum* (Pt), *Pardosa amentata* (Pa), *Pardosa pseudoannulata* (Pan), *Phalangium opilio* (Po), *Pholcus phalangioides* (Pp), *Phoxichildium femoratum* (Pf), *Stegodyphus dumicola* (Sd), *Strigamia maritima* (Sm), *Tetranychus urticae* (Tu), and *Tribolium castaneum* (Tc). Node labels indicate ultrafast bootstrap support values. See Supplementary File 1 for accession numbers, Supplementary File 9 for amino acid sequence alignments, and Supplementary File 10 for nucleotide sequence alignments of *Wnt1/wg* duplicates in *C. acosta, C. sculpturatus* and *Euphrynichus bacillifer*.

Duplicates of *Wnt11* previously identified in *P. tepidariorum* (Janssen et al. 2010) were recovered from all surveyed spiders except *A. geniculata*, *C. acosta*, and both scorpions. The spider Wnt11 duplicates formed two separate and well-supported clades (≥85%, Figure 4), and each amblypygid Wnt11 orthology group was sister to one of the spider clades (98-99%; Figure 4). Similarity between paralogs in the four new transcriptomes was very low (41-54%). We propose that the spider and amblypygid Wnt11 duplicates are probably retained from the ancestral WGD. The resolution of the scorpion Wnt11 duplicates is less clear; ortholog pairs resolve together with 100% support, but only one resolves as sister to the two clear clades occupied by the spider and amblypygid duplicates (98%; Figure 4). They exhibit similar sequence similarity (49-50%) to the putative ohnologs.

Wnt4 paralogs from *M. muscosa, P. amentata, A. geniculata, P. pseudoannulata* and *S. dumicola* form two separate and well-supported clades with duplicates from the amblypygids (bootstrap ≥ 91%). They show substantial sequence divergence within species (53-64% similarity), indicating that they are again likely to represent retained ohnologs following the arachnopulmonate WGD, despite being lost in the lineage to *P. tepidariorum*.

We did not recover duplicates of *Wnt2*, *Wnt5*, *Wnt8-10*, or *Wnt16* in any arachnopulmonate lineage. This suggests their loss shortly after WGD in the common ancestor of all arachnopulmonates. Two dissimilar overlapping partial sequences of *WntA* were recovered from the genome of *P. pseudoannulata*, but these resolve as sister to one another (Figure 4) and are located on the same scaffold, indicating a lineage-specific tandem duplication. We therefore suggest that the *WntA* duplicate resulting from WGD was also lost in the spider ancestor.

We identified two copies of *Wnt1/wg* in the transcriptomes of both amblypygids and *A. geniculata*, and in the genomes of *P. pseudoannulata, C. sculpturatus*, and *M. martensii*. To the best of our knowledge, this is the first duplication of *Wnt1/wg* reported in any animal. This requires critical interpretation. We can eliminate the possibility of individual variation in *C. acosta,* as embryos are produced by parthenogenesis and are therefore clones, and in *C. sculpturatus* and *S. dumicola,* as the sequences were recovered from a single individual’s genome (Supplementary File 1). Sequence similarity between paralogs was low (55-73%) compared to similarity between orthologs at the order level (e.g. 91% between *M. muscosa* and *P. amentata*), and comparable to orthologs at the class level (e.g. 61% between *P. tepidariorum* and *Ixodes scapularis*), reducing the likelihood that we are detecting allelic variation within individuals. We also inspected nucleotide alignments and found lower paralog sequence similarity than evident from amino acid sequences (65-69%), indicating synonymous evolution. Although synteny analysis is required for conclusive confirmation, our phylogeny indicates that the amblypygid, scorpion, and *A. geniculata* duplicates are likely to be ohnologs retained from the arachnopulmonate WGD, as they form separate clades (≥70%, Figure 4). The putative duplicates in *P. pseudoannulata*, in contrast, resolve as sister to one another. These two sequences have (short) non-identical overlapping regions, but they are partial. It is equivocal whether they represent a genuine, lineage-specific duplication.

### Frizzled repertoires and their origins

All surveyed spiders possess at least one copy of FzI and FzII, with a second copy recovered from the genome of *Stegodyphus dumicola* and the transcriptome of *Marpissa muscosa*, respectively (Figure 5). FzIII was absent from the transcriptomes of *M. muscosa* and *P. amentata,* and the genomes of *S. dumicola* and *P. pseudoannulata*, consistent with *P. tepidariorum* (Janssen et al. 2015). However, single FzIII orthologs were recovered from *Acanthoscurria geniculata* and previously identified in *Pholcus phalangioides* (Janssen et al. 2015). Thus, entelegynes may universally lack *fz3* but it was likely present in the ancestor of all spiders. Consistent with *P. tepidariorum* (Janssen et al. 2015), we identified two FzIV sequences in *S. dumicola* and *A. geniculata*, but only one in *M.* muscosa, *P. amentata* and *P. pseudoannulata* and only one was previously recovered from *P. phalangioides* (Janssen et al. 2015). Janssen et al. (2015) demonstrated that the expression of the two *fz4* paralogs in *P. tepidariorum* is separated temporally, so we might not expect to detect both transcripts in developing embryos of similar stages. However, this does not explain the absence of a FzIV duplicate from the genome of *P. pseudoannulata*.

**Figure 5.**
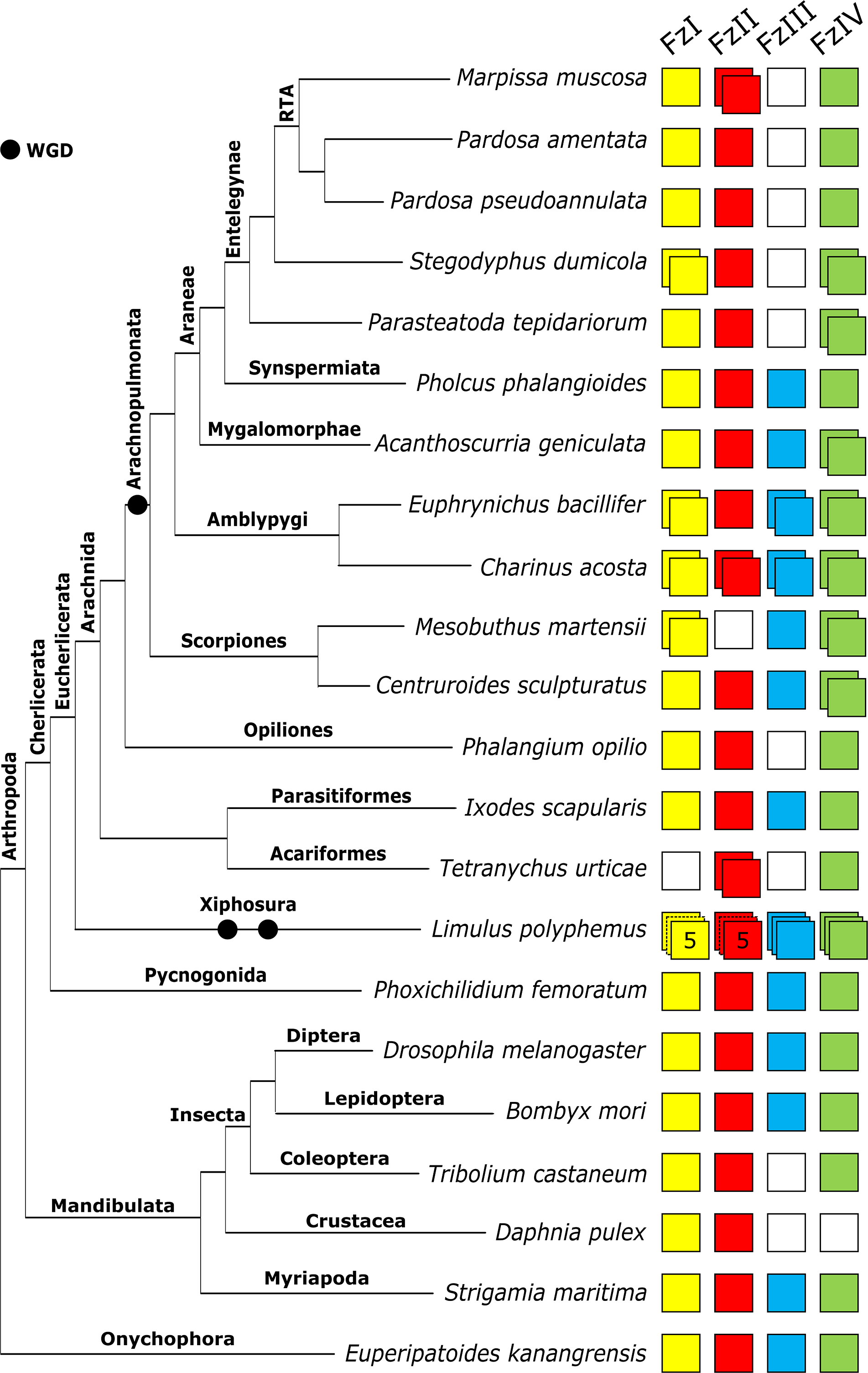
Repertoire of *frizzled* genes in arachnids and other selected arthropods. The four *frizzled* orthology groups (FzI, FzII, FzIII, and FzIV) are represented by coloured boxes, with duplicated genes represented by overlapping boxes and gene loss represented by a white box. Figure includes *frizzled* repertoires previously surveyed in the arachnids *Parasteatoda tepidariorum, Mesobuthus martensii*, *Ixodes scapularis* (all genomes), and *Pholcus phalangioides* (embryonic transcriptome); the myriapod *Strigamia maritima*; the insects *Drosophila melanogaster* and *Tribolium castaneum*; and the onychophoran *Euperipatoides kanangrensis*.

Both amblypygid species have a large repertoire of *frizzled* genes compared to other arachnids, with duplicates of FzI, FIII and FzIV in both species and an additional duplicate of FzII in *Charinus acosta* (Figures 5).

The scorpions *Mesobuthus martensii* (Janssen et al. 2015) and *Centruroides sculpturatus* possess single FzI and FzIII orthologs and a duplication of FzIV. We also recovered a FzII ortholog in *C. sculpturatus*, and a FzI duplicate in *M. martensii*.

Among the non-arachnopulmonate arachnids, *Ixodes scapularis* possesses a single copy of all four orthology groups, but FzIII was absent in *Phalangium opilio* and *Tetranychus urticae*. A FzII duplicate was recovered from *Tetranychus urticae. Phoxichilidium femoratum* and *Limulus polyphemus* possess all four orthology groups. No duplicates were recovered in *P. femoratum*. Five copies of FzI and FzII and three copies of FzIII and FzIV were recovered from *L. polyphemus* (Figure 5).

Frizzled paralogs appear to stem from both WGD events and lineage-specific duplications. FzI duplicates were recovered from both amblypygid transcriptomes*, M. martensii*, and *S. dumicola.* The Sd-Fz1 and Me-Fz1 paralog pairs exhibit high sequence similarity (>73%) and resolved as sisters with 100% bootstrap support, indicating that these are likely the results of lineage-specific duplications. Conversely, the amblypygid Fz1-1 and Fz1-2 ortholog pairs form separate clades with the Fz1 sequences from spiders and scorpions, respectively (Figure 6). We therefore interpret these duplicates as ohnologs.

**Figure 6.**
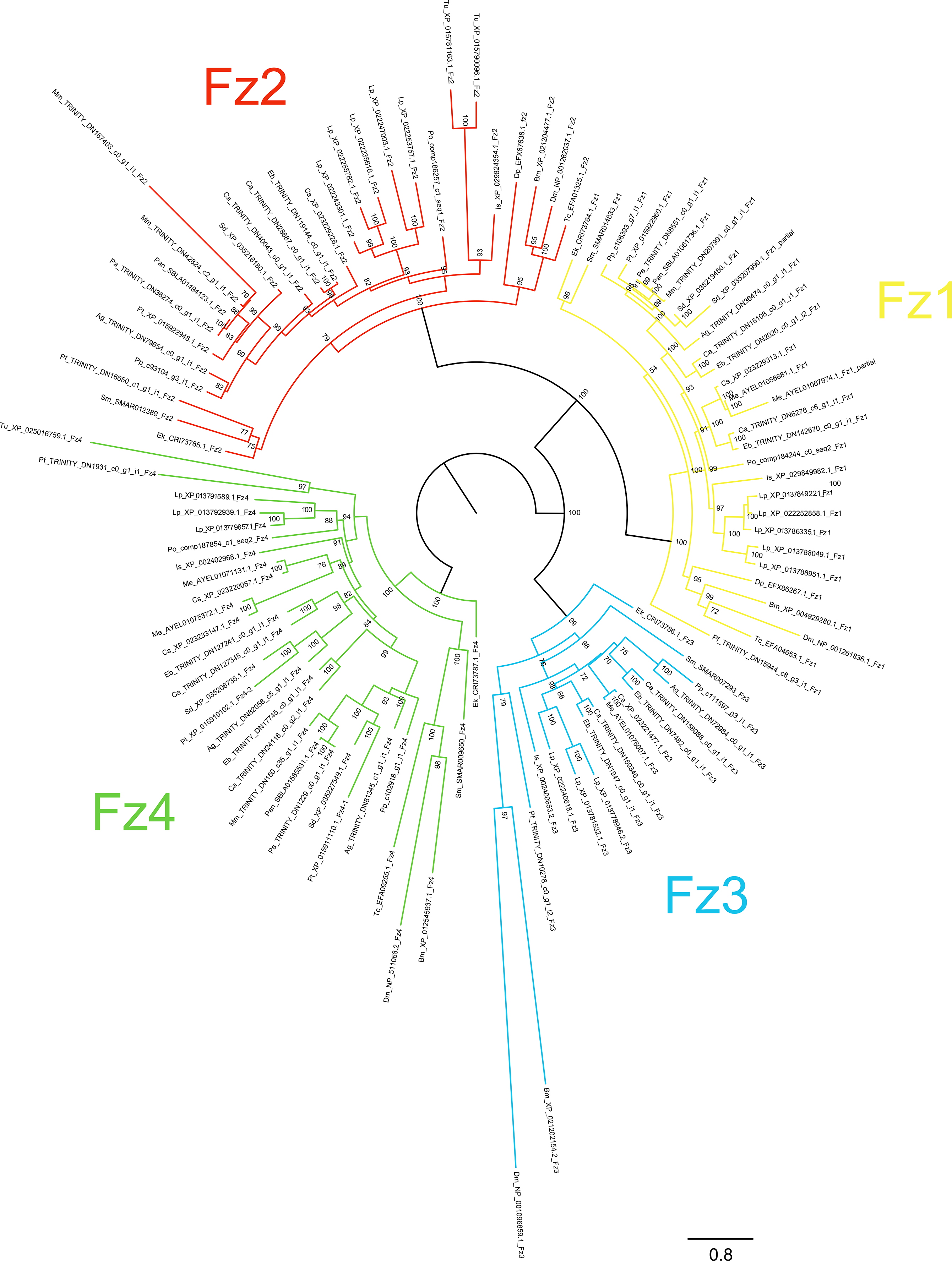
Maximum likelihood phylogeny of Frizzled proteins. The *frizzled* genes are shown as different colours (after Figure 5). Panarthropods included: *Acanthoscurria geniculata* (Ag), *Bombyx mori* (Bm), *Centruroides sculpturatus* (Cs), *Charinus acosta* (Ca), *Drosophila melanogaster* (Dm), *Euperipatoides kanangrensis* (Ek), *Euphrynichus bacillifer* (Eb), *Ixodes scapularis* (Is), *Limulus polyphemus* (Lp), *Marpissa muscosa* (Mm), *Mesobuthus martensii* (Me), *Parasteatoda tepidariorum* (Pt), *Pardosa amentata* (Pa), *Pardosa pseudoannulata* (Pan), *Phalangium opilio* (Po), *Pholcus phalangioides* (Pp), *Phoxichildium femoratum* (Pf), *Stegodyphus dumicola* (Sd), *Strigamia maritima* (Sm), *Tetranychus urticae* (Tu), and *Tribolium castaneum* (Tc). Node labels indicate ultrafast bootstrap support values. See Supplementary File1 for accession numbers and Supplementary File 11 for alignments

Duplicates of FzII were recovered from *M. muscosa* and *C. acosta*. The Mm-Fz2 paralogs form a well-supported clade (79%; Figure 6), indicating that this is the result of a lineage-specific duplication followed by rapid sequence divergence in Mm-Fz2-2 (53% sequence similarity). The origin of the FzII duplication in *Charinus acosta* is not clear. Ca-Fz2-1 resolves as a sister group to the spider Fz2 sequences (99%; Figure 6) and Ca-Fz2-2 forms a clade with Eb-Fz2, which in turn resolves as sister to the Ca-Fz2-1 and spider sequences (93%; Figure 6). This topology could support an ohnolog relationship between Ca-Fz2-1 and Ca-Fz2-2 but cannot be confirmed. Sequence similarity between the *C. acosta* paralogs is relatively high (82%), perhaps higher than expected from ohnologs.

The origin of the amblypygid FzIII duplicates is also unclear. One ortholog pair forms a clade with *L. polyphemus* Fz3 (66%; Figure 6) and the other forms a clade with the spiders and scorpions (≥ 70%; Figure 6). This suggests an origin in WGD, but support for their placement is not strong (66%, Figure 6). Paralogous pairs demonstrate middling sequence similarity (65-66%).

Both amblypygids and scorpions, and the spiders *P. tepidariorum, S. dumicola* and *A. geniculata*, possess FzIV duplicates. The spider Fz4 sequences formed two separate and well-supported clades (100%; Figure 6). The amblypygid ortholog pairs resolved as sister to the two spider clades (98%; Figure 6). The scorpion ortholog pairs, however, formed a clade together (76% support; Figure 6) which is sister to all the spider and amblypygid Fz4 sequences (82%; Figure 6). All paralogous pairs exhibited substantial sequence divergence (similarity 44-58%). We propose that the spider and amblypygid FzIV duplicates are retained from the ancestral WGD; however, once again, the origin of the scorpion FzIV duplicates is less clear, and may reflect a lineage-specific duplication.

## DISCUSSION

The impact of WGD on genome evolution is evident across the three gene families surveyed here, but it appears that retention patterns of putative ohnologs vary substantially between them. We also see distinct phylogenetic patterns beginning to emerge in gene repertoires, with improved sampling within lineages enabling us to distinguish between possible signal and likely noise. Of course, transcriptomic data come with necessary caveats regarding gene expression and capture, and absences from transcriptomes should be regarded with caution. However, where patterns are consistent across multiple species, or better, between transcriptomic and genomic resources, we can be more confident about their authenticity. For example, comparisons of the transcriptome of *Pardosa amentata* and the genome of *Pardosa pseudoannulata* show identical Hox and Fz repertoires, indicating good gene capture in the former. The apparent duplicates of *Wnt1/wg* and *WntA* in *P. pseudoannulata* that were not recovered in *P. amentata* are of uncertain status (partial sequences, phylogenetic position) and were absent from all other spider genomes.

### Hox duplicates are broadly retained, with slight variation between major clades

The widespread retention of duplicate Hox genes is consistent among the arachnopulmonate orders studied to date, and specific repertoires appear to be fairly conserved at the order level (Schwager et al. 2007; Cao et al. 2013; Di et al. 2015; Schwager et al. 2017; Leite et al. 2018; Ontano et al. 2020). Given that this is the level at which overall body plans are conserved, this is perhaps not surprising. Thanks to the relatively conserved expression patterns of Hox genes along the antero-posterior axis of chelicerates, we can begin to speculate about the possible macroevolutionary implications of duplication and loss. For example, an anticipated duplicate of *pb* appears to have been lost in both amblypygids but persists in spiders and scorpions. In spiders, both *pb* paralogs are expressed in the pedipalp and leg-bearing segments, separated temporally (Schwager et al. 2017). Given the highly derived nature of the raptorial pedipalps and the antenniform first pair of walking legs in amblypygids, it is perhaps surprising that this duplicate was not retained. However, this might indicate that other Hox genes expressed in the anterior prosomal segments (e.g. *lab*, *Hox3*, or *Dfd*) may contribute to these morphological innovations. Recent work by Gainett and Sharma (Gainett and Sharma 2020) examining the specification of the antenniform legs found little difference in *Distal-less*, *dachshund* or *homothorax* expression between that and posterior leg pairs, indicating that these are not likely to be responsible. A good candidate for future study might be *lab*: a single ortholog is expressed in both the pedipalps and the first walking leg in the harvestman *Phalangium opilio* (Sharma et al. 2012), and expression patterns and experimental manipulation provide evidence for functional divergence between the two *lab* paralogs, also expressed in the pedipalps and first walking legs, in *P. tepidariorum* (Pechmann et al. 2015; Schomburg et al. 2020).

Likewise, the seemingly universal absence of a *ftz* duplicate in spiders, but its retention in scorpions and amblypygids, poses a similar question. In both *P. opilio* (Sharma et al. 2012) and *P. tepidariorum* (Schwager et al. 2017), a single copy of *ftz* is expressed in leg pairs 2-4, implying that spiders have not lost *ftz* functionality by losing a duplicate. Any subfunctionalization or neofunctionalization that could be evident in scorpion and amblypygid *ftz* had therefore presumably not taken place at the point of their divergence from spiders. Expression patterns of the two paralogs in these groups would be of interest for comparison with both harvestmen and spiders. Given that neither scorpions nor amblypygids exhibit much divergence within L2-4, there are no obvious routes for subfunctionalization.

The potential loss of a *Hox3* duplicate in the spider RTA clade would be an unusual example of infraorder variation in Hox repertoires, if it represents a true absence. The consistency of this finding between both transcriptomic and genomic resources supports this conclusion. In *P. tepidariorum*, Hox3-A and Hox3-B exhibit dramatic protein sequence divergence (20.6% similarity). Both copies are expressed in the embryo, with their expression overlapping spatially and temporally but not identical, indicating some functional divergence; however, *Hox3-A* expression was reported to be very weak (Schwager et al. 2017). In the other spiders for which we recovered *Hox3* duplicates, paralogs also exhibited very low sequence similarities (24.7-30.6%), and these duplicates had very low similarity to each other (17.2-30.1%, compared to 33-58.2% between those resolving with other spider Hox3 sequences). In all cases, one resolved within a well-supported clade of spider Hox3 and the other was placed haphazardly. We speculate that the *Hox3* duplicate is highly divergent or degenerate in spiders, leading to eventual pseudogenization in the RTA clade. As other infraorder losses of Hox genes were only observed in single species, some from embryonic transcriptomes, it would be premature to conclude that they are genuinely absent from the genome.

### The retention of Wnt duplicates is more specific, but with some surprises

In contrast to the widespread retention of Hox duplicates, these new data indicate that the retention of duplicate Wnt genes is less common and restricted to certain subfamilies. Apparent ohnologs of *Wnt4*, *Wnt7* and *Wnt11* are retained in the majority of arachnopulmonates, for example, but *Wnt2*, *Wnt5*, *Wnt8-10*, and *Wnt16* are present in single copies across the board. These patterns could reflect early losses, by chance, of duplicates, or differential ‘evolvability’ of Wnt subfamilies. The fact that retention patterns appear to differ between the arachnopulmonate and horseshoe crab independent WGD events lends support to the former hypothesis.

Our understanding of specific Wnt functions among arthropods is more limited than that of Hox genes, but Wnt expression patterns in *Parasteatoda tepidariorum* are available for tentative comparison. For example, previous attempts to characterise the expression patterns of *Wnt11* paralogs in *P. tepidariorum* only detected expression of *Wnt11-2* (Janssen et al. 2010). Given the retention of *Wnt11-1* in both spiders and scorpions, and the considerable divergence between paralogous sequences, *Wnt11* could be a good candidate for sub- or neofunctionalization, but the role of *Wnt11-1* remains unknown. Conversely, the presence of two apparent *Wnt4* ohnologs in spiders and amblypygids (Figures 3-4) contrasts with the retention of only one *Wnt4* in *P. tepidariorum*. The role of the additional copy, particularly in spiders, will be of future interest and expression patterns of *Wnt4-2* in these groups will help to clarify this. Although it is possible that the second copy is redundant or in the process of pseudogenization, the fact that both ohnologs are retained in two large clades, with detectable levels of expression during development, suggests that this is not the case. Thus, its absence in *P. tepidariorum* unexpectedly appears to be the exception.

The discovery of duplicate *Wnt1/wg* in scorpions, amblypygids and mygalomorphs is particularly exciting: duplicates of this Wnt gene have not yet been detected in any other metazoans, even following multiple rounds of WGD in vertebrates, teleosts (see https://web.stanford.edu/group/nusselab/cgi-bin/wnt/vertebrate), or horseshoe crabs. Wnt1/wg performs a wide variety of roles in arthropods, including in segment polarization and in appendage and nervous system development (Murat et al. 2010) and has an accordingly complex expression pattern in *P. tepidariorum*, appearing in the L1 and L2 segments, limb buds, and dorsal O2 and O3 segments (Janssen et al. 2010). In theory, therefore, there is ample potential for subfunctionalization. Gene expression and functional studies of *Wnt1/wg* duplicates in arachnopulmonates will no doubt prove extremely interesting in the future.

The presence of *Wnt10* in both amblypygids and *Phalangium opilio* is also intriguing because it is absent from all other chelicerates surveyed so far. Whether this indicates multiple losses of *Wnt10* in all other arachnid lineages, the recovery of a lost *Wnt10* in amblypygids and harvestmen, or the co-option of another gene, is unclear. It is notable, however, that *Wnt10* is also absent in both *Limulus polyphemus* and *Phoxichilidium femoratum*, non-arachnid chelicerates. This might favour the recovery or co-option hypotheses.

### Fz repertoires vary substantially

Previous studies of spiders, scorpions, and ticks indicated that Frizzled repertoires in these groups are restricted to three or four copies, often with incomplete representation of the four orthology groups (Janssen et al. 2015). The spiders *Marpissa muscosa*, *Stegodyphus dumicola*, *Pardosa pseudoannulata* and *Pardosa amentata* are consistent with this pattern, albeit with a unique duplication of FzII in the jumping spider. We also recovered a single copy of FzII in *Centruroides sculpturatus*, which is absent in *Mesobuthus martensii*, and a FzI duplicate in *M. martensii* that was previously missed. In contrast, all four frizzled subfamilies were recovered in both amblypygid species, with three present in duplicate in *Euphrynichus bacillifer* and four in *Charinus acosta*. Based on our data, it appears that the frizzled repertoire of amblypygids is around twice the size of all other arachnids and may have followed a very different evolutionary trajectory to spiders and scorpions following WGD. The expanded repertoire of frizzled genes in amblypygids is intriguing since they have the widest Wnt repertoires, via both duplication and representation of the subfamilies (Wu and Nusse 2002). However, although frizzled genes encode key receptors for Wnt ligands, they have other Wnt-independent functions, so the expansion of the frizzled gene repertoire could be equally related to the evolution of alternative signalling roles (Janssen et al. 2015; Yu et al. 2020).

### Arachnopulmonate genome evolution in the wake of WGD

Our new analyses provide a thorough survey of Hox, Wnt and frizzled genes in arachnids, and substantially improve the density and breadth of taxonomic sampling for these key developmental genes in Arachnopulmonata. We find evidence of consistent evolutionary trajectories in Hox and Wnt gene repertoires across three of the five arachnopulmonate orders, with inter-order variation in the retention of specific paralogs. We have also identified intraorder variation at the level of major clades in spiders, which could help us better understand their morphological evolution. In new data for a third arachnopulmonate lineage, the amblypygids, we find additional evidence supporting an ancestral WGD and are better able to reconstruct the chronology of gene duplications and losses in spiders and scorpions. These transcriptomic resources are among the first available for amblypygids and will aid future investigations of this fascinating group.

By improving taxonomic coverage within the spider lineage we are better able to polarise some loss/duplication events and identify potential new trends within the spiders, particularly illustrating separations between synspermiatan and entelegyne spiders, and between the derived RTA clade and other spiders. Despite being unable to ultimately conclude that some missing transcripts reflect genuine genomic losses, it appears that the evolution of these developmental genes in spiders is more complicated than we thought. It may be that these gene repertoires are genuinely more variable within spiders than they are in amblypygids or scorpions; spiders are by far the most taxonomically diverse arachnopulmonate order, and the apparent diversity of repertoires may simply reflect this. Conversely, the higher apparent intraorder diversity of gene repertoires may be an artefact of increased sampling in spiders (up to four or five species for specific gene families) compared to the one or two available resources for scorpions and amblypygids; we may detect more diversity within these groups with increased sampling. Nonetheless, we see two notable trends within spiders, outlined below.

First, we see several characters that appear to unite the RTA clade, which contains almost half of all extant spider species (World Spider Catalog 2019), having diversified rapidly following its divergence from the orb weavers (Garrison et al. 2016; Fernández et al. 2018; Shao and Li 2018). *Marpissa muscosa, Pardosa amentata,* and *Pardosa pseudoannulata* all exhibit the apparent loss of *Hox3* and *fz4* paralogs and the retention of a *Wnt4* duplicate. The identification of genetic trends potentially uniting this group is exciting, even if the macroevolutionary implications are unclear: as described above, the possible functions of a *Wnt4* paralog are elusive. Members of the RTA clade are very derived compared to other araneomorph spiders, both morphologically (e.g. male pedipalp morphology and sophisticated eyes) and ecologically (most are wandering hunters), and their rapid diversification would align with clade-specific genetic divergence (Garrison et al. 2016; Fernández et al. 2018; Shao and Li 2018).

Second, although data are only available for single representatives of the plesiomorphic clades Synspermiata (*Pholcus phalangioides*) and Mygalomorphae (*Acanthoscurria geniculata*), these hint at lineage-specific losses of Hox paralogs and recover the only examples of FzIII found in spiders so far. The presence of FzIII is consistent with other arachnopulmonate groups and suggests that it was present in the spider ancestor and only lost in the more derived entelegyne lineages. If selected Hox duplicates are indeed absent from the genomes of these two species, this represents an interesting divergence between the three major groups of spiders. Although it may seem unusual for Hox duplicates to be lost within lineages, a recent high-quality genome of *Trichonephila antipodiana* revealed likely absences of *Hox3*, *Ubx* and *abdA* ohnologs in addition to *ftz* (Fan et al. 2021). Though they are unlikely to be directly responsible, the apparent divergence in gene repertoires we see between *A. geniculata*, *P. phalangioides* and the other spider lineages might provide a starting point for understanding the important morphological differences between mygalomorphs, synspermiatans and entelegynes. However, genomic information for additional taxa in both groups is required to verify these potential losses.

The amblypygids emerge as a key group of interest for studying the impacts of WGD owing to their high levels of ohnolog retention. Our transcriptomes, from representatives of two major clades, provide new evidence supporting a WGD in the ancestor of arachnopulmonates and demonstrating widespread retention of ohnologs in three major families of developmental genes (consistent with the retention of many duplicated regulators of eye development in other species, Gainett et al. 2020). In all three gene families we studied, repertoires were largest in the amblypygid species. This was particularly the case in *Charinus acosta*, which belongs to the less speciose and more plesiomorphic infraorder Charinidae within living Amblypygi. Although this study represents just two amblypygid species and three gene families, this appears to contradict widespread predictions of diversification with the duplication of important developmental genes such as Hox (e.g. Van De Peer et al. 2009). Of particular interest are the amblypygid Wnt gene repertoires. We have identified from their transcriptomes, and from the published genome of *Centruroides sculpturatus*, the first reported duplicates of *Wnt1/wg* in any animal, as well as the first reported *Wnt10* in any arachnid. Future functional studies of these genes and their expression during development will be critical to understanding the evolutionary impacts of these unusual components of amblypygid gene repertoires. Amblypygids also represent a potential model group for studying the evolution of arthropod body plans, owing to the unusual and derived morphology of the pedipalps and especially the first walking legs. Thanks to a substantial existing body of work on anterior-posterior patterning, segmentation and appendage development in spiders and other arachnids, we may have a chance to crack the genetic underpinnings of these dramatic evolutionary innovations (Pechmann et al. 2009; Sharma et al. 2012; Sharma et al. 2014; Turetzek et al. 2016; Schwager et al. 2017; Turetzek et al. 2017; Schomburg et al. 2020; Baudouin-Gonzalez et al. 2021).

Finally, our analysis of existing genomic data for *Centruroides sculpturatus* and *Mesobuthus martensii* has recovered several Wnt and Frizzled gene duplications, similar to spiders and amblypygids. However, in contrast to those groups, our phylogenies have sometimes supported within-lineage duplication in scorpions, as opposed to the retention of ohnologs following WGD, even when these are observed in spiders and amblypygids. This was the case for *AbdB, Wnt1/wg, Wnt6, Wnt7,* and FzIV (Figures 2,4,6). However, levels of sequence similarity in these cases were comparable for *C. sculpturatus* paralogs and amblypygid and spider ohnologs, when we might expect within-lineage duplicates to show higher similarity. The resolution of the paralogous sequences in our phylogenetic analyses could be confounded by the early-branching position of scorpions within Arachnopulmonata, which means paralogs would be expected to appear towards the bottom of ortholog clades and are more vulnerable to movement. Nonetheless, this pattern emerged multiple times in our analyses and may be of future interest.

Both arachnopulmonates and horseshoe crabs have been subject to WGD. Comparison between these independent events is a useful tool in studying WGD, spanning smaller phylogenetic distances than arachnopulmonate-vertebrate comparisons. From the three gene families surveyed here, both patterns and inconsistencies emerge. As in arachnopulmonates, Hox gene duplicates appear to be overwhelmingly retained, even in triplet or quadruplet in some cases, in *Limulus polyphemus*. *Ftz* is an interesting exception that aligns with the absence of duplicates in spiders, but not scorpions or amblypygids. However, the apparent loss of all *Ubx* and two *abdA* duplicates stands at odds with the arachnopulmonates, wherein these are largely retained. Wnt repertoires follow a similar pattern to arachnopulmonates - only select genes are retained in duplicate - but its application is very different: for example, whereas *Wnt4* duplicates are common in arachnopulmonates, they are completely absent in *L. polyphemus*, and vice versa for *Wnt5*. There are, however, some Wnts that are commonly or even universally present in single copies following both independent WGD events, potentially indicating low potential for sub- or neofunctionalization, such as *Wnt8*, *Wnt6* and *Wnt1*/*wg*.

Overall, our new data provide further evidence of an ancestral arachnopulmonate WGD, identify evolutionary patterns within gene families following WGD, reveal new diversity in spider gene repertoires, better contextualise existing data from spiders and scorpions, and broaden the phylogenetic scope of available data for future researchers. However, other arachnid groups, both with and without ancestral WGD, require further study. Recent work on two pseudoscorpions recovered duplicates of most Hox genes (Ontano et al. 2020), in contrast to previous surveys (Leite et al. 2018). This not only further supports arachnopulmonate WGD but substantially improves our understanding of pseudoscorpion placement within arachnids, which has been historically problematic. The sequencing of a pseudoscorpion genome provides the tantalising chance to add a fourth lineage to future studies of WGD and its impacts (Ontano et al. 2020). The remaining arachnopulmonate orders, thlyphonids (vinegaroons or whip scorpions) and schizomids, form a clade with amblypygids (Pedipalpi) and should also have been subject to the arachnopulmonate WGD. Future work on these groups will shed light on the unusual patterns of gene retention we find in both major clades of amblypygids. Non-arachnopulmonate arachnids are invaluable for contextualising the changes that occur following WGD, both in terms of gene repertoires and of gene function. Genomic resources and gene expression pattern studies are vital for this, and the harvestman *Phalangium opilio* has emerged as the clear model species (Sharma et al. 2012; Gainett et al. 2021). The ability to compare rates of sequence divergence, within-lineage gene duplication, and, eventually, functional properties of developmental genes in these groups will provide critical comparative data for arachnopulmonates.

## Availability of data and materials

Alignments and gene accession numbers are included as supplementary information files. Transcriptomes are available via SRA (BioProject PRJNA707377). Assemblies will be made publicly available upon acceptance for publication.

## Competing interests

The authors declare that they have no competing interests.

## Funding

This work was supported by the John Fell Fund, University of Oxford (award 0005632 to LSR), NERC (NE/T006854/1 to APM and LSR), the Leverhulme Trust (RPG-2016-234 and RPG-2020-237 to APM and LSR, respectively), and the Biotechnology and Biological Sciences Research Council (BBSRC) (grant number BB/M011224/1). These funding bodies were not involved in the design of this study, data collection or analysis, or in writing the manuscript.

## Authors’ contributions

AH analysed assembled transcriptomes, identified gene candidates, performed phylogenetic analyses, interpreted data, prepared figures and wrote sections of the manuscript. LBG contributed to data analysis and interpretation. AS processed embryos and performed RNA extractions. RJ provided data for *Pholcus phalangioides* and *Phalangium opilio*. MS provided amblypygid embryos. MH provided sequence data and contributed to phylogenetic analysis. SA assembled transcriptomes, analysed assembly quality and wrote sections of the manuscript. APM designed and supervised the project, contributed to data interpretation and analysis, and wrote sections of the manuscript. LSR designed and supervised the project, performed phylogenetic analyses, interpreted data, prepared figures and wrote the manuscript. All authors read and approved the final manuscript.

## Acknowledgements

The authors are very grateful to Philip Steinhoff and Gabriele Uhl (University of Greifswald) for providing embryos of *Marpissa muscosa*, to Matthias Pechmann (University of Köln), Natascha Turetzek (Ludwig-Maximilians-University of Munich) and Prashant Sharma (University of Wisconsin Madison) for providing assembled transcriptomes of *Acanthoscurria geniculata*, *Pholcus phalangioides* and *Phalangium opilio*, respectively, to Nathan Kenny for helpful comments on the manuscript, and to Simon Ellis (University of Oxford) for IT support.

## Notes

### Competing Interest Statement

The authors have declared no competing interest.

### Summary of Updates

- Expansion of taxa surveyed, to include 16 chelicerate species - Balancing of taxa between gene families - Addition of phylogenetic analyses of Hox homeodomain sequences in RAxML and Iqtree

