## Supplementary figures and images for "Widespread retention of ohnologs in key developmental gene families following whole genome duplication in arachnopulmonates"

### SF5_Hox_prot_HD.iqtree.pdf

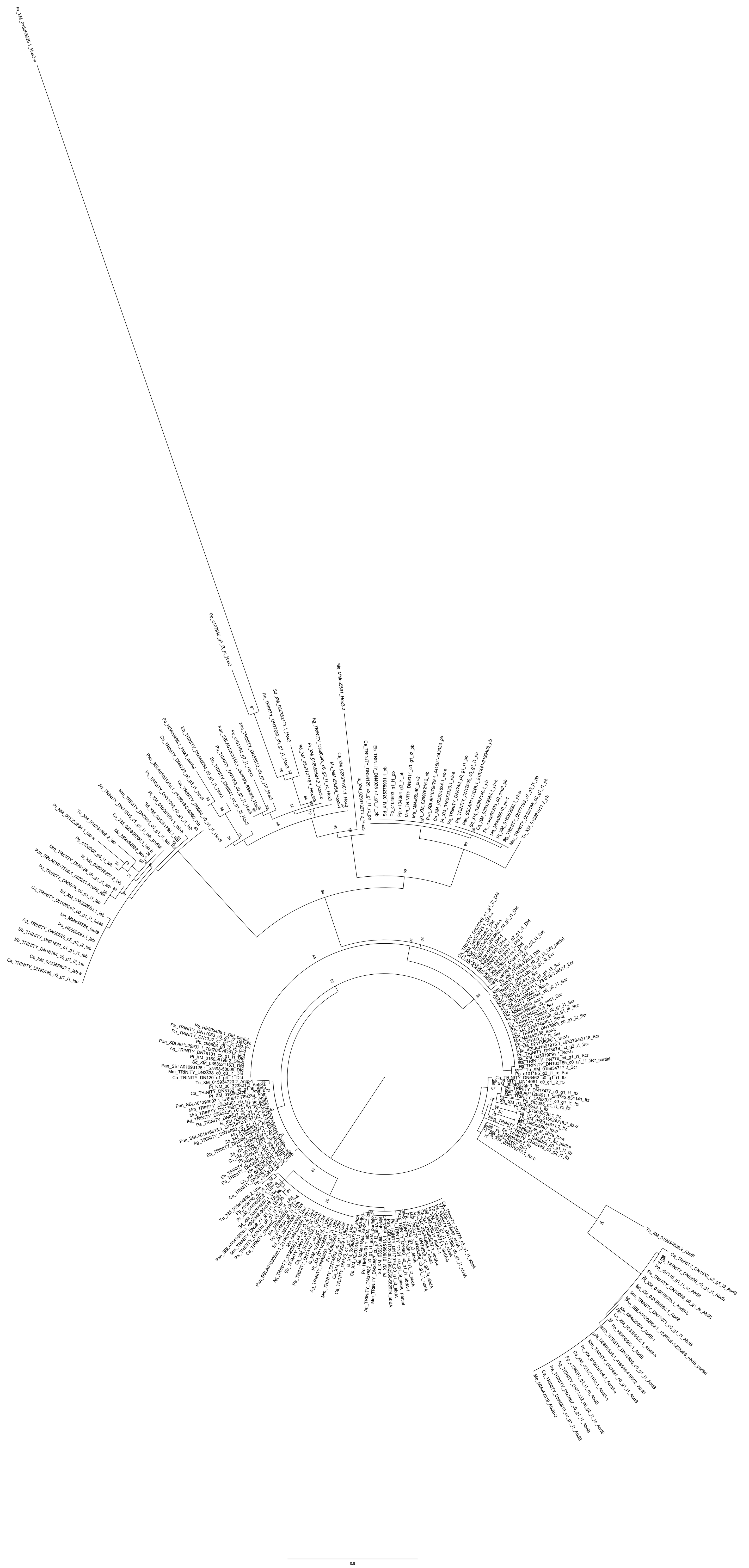

### SF6_Hox_prot_HD.raxml.pdf

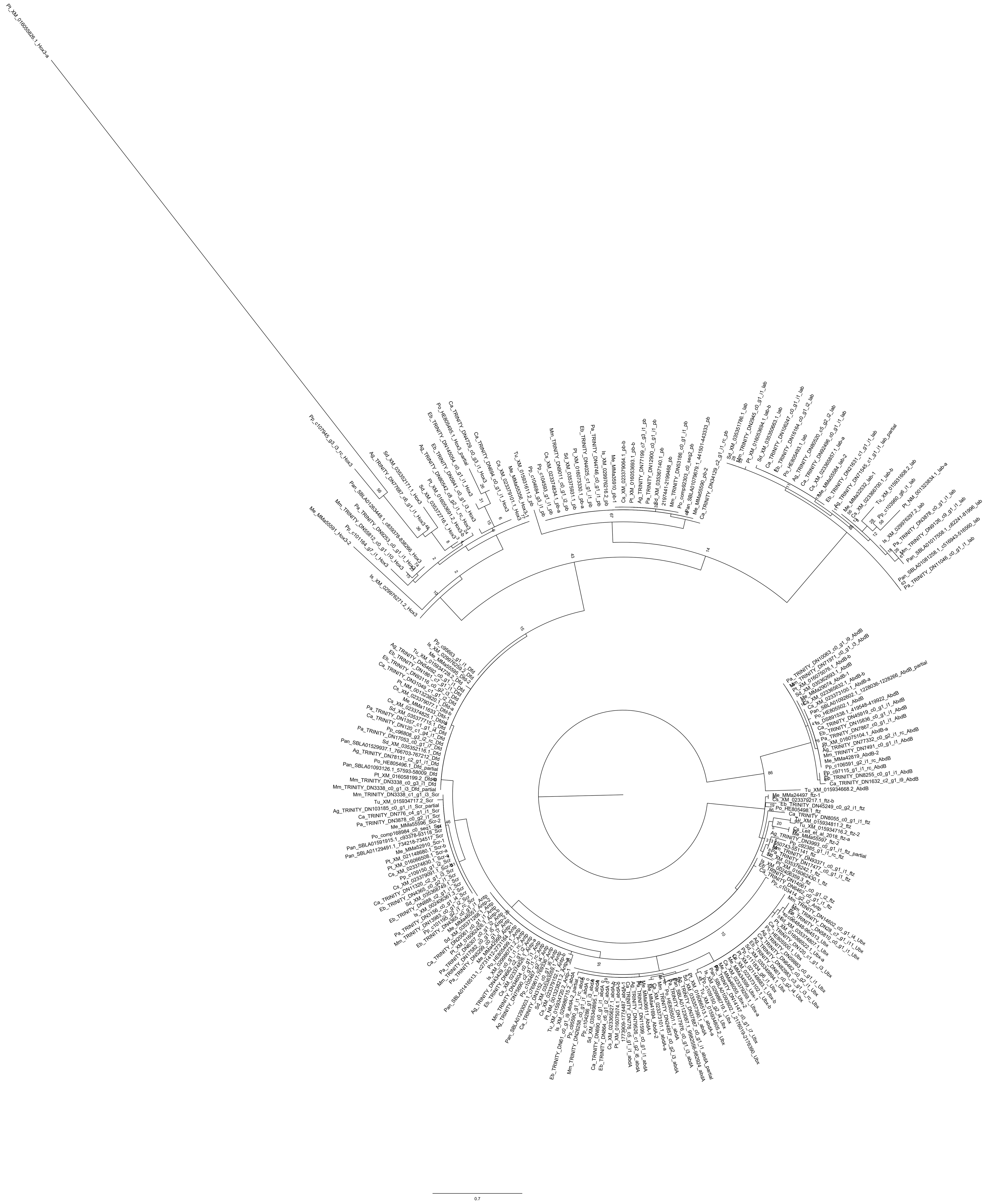

### SF7_Hox_gene_HD.iqtree.pdf

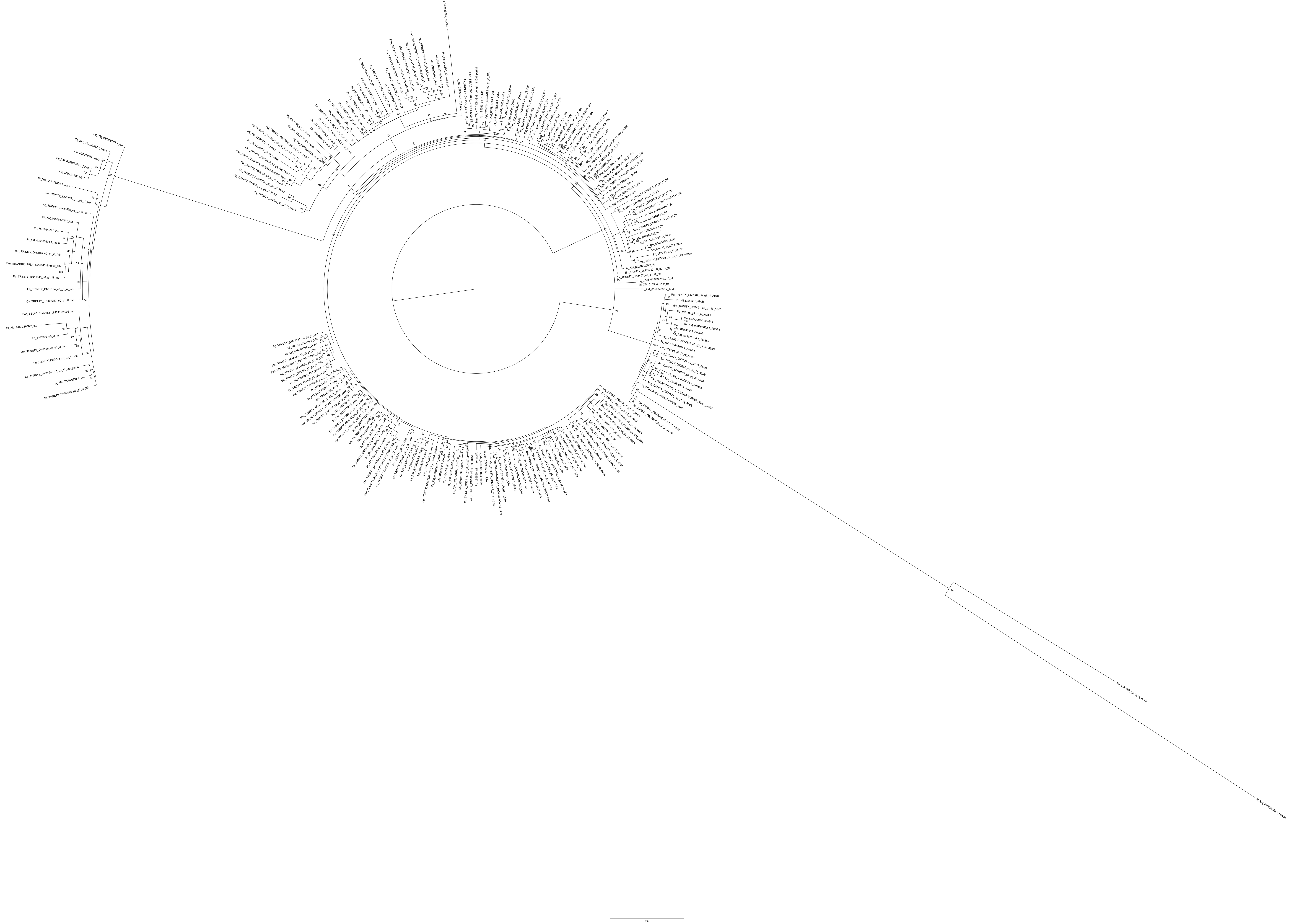

### SF8_Hox_gene_HD.raxml.pdf

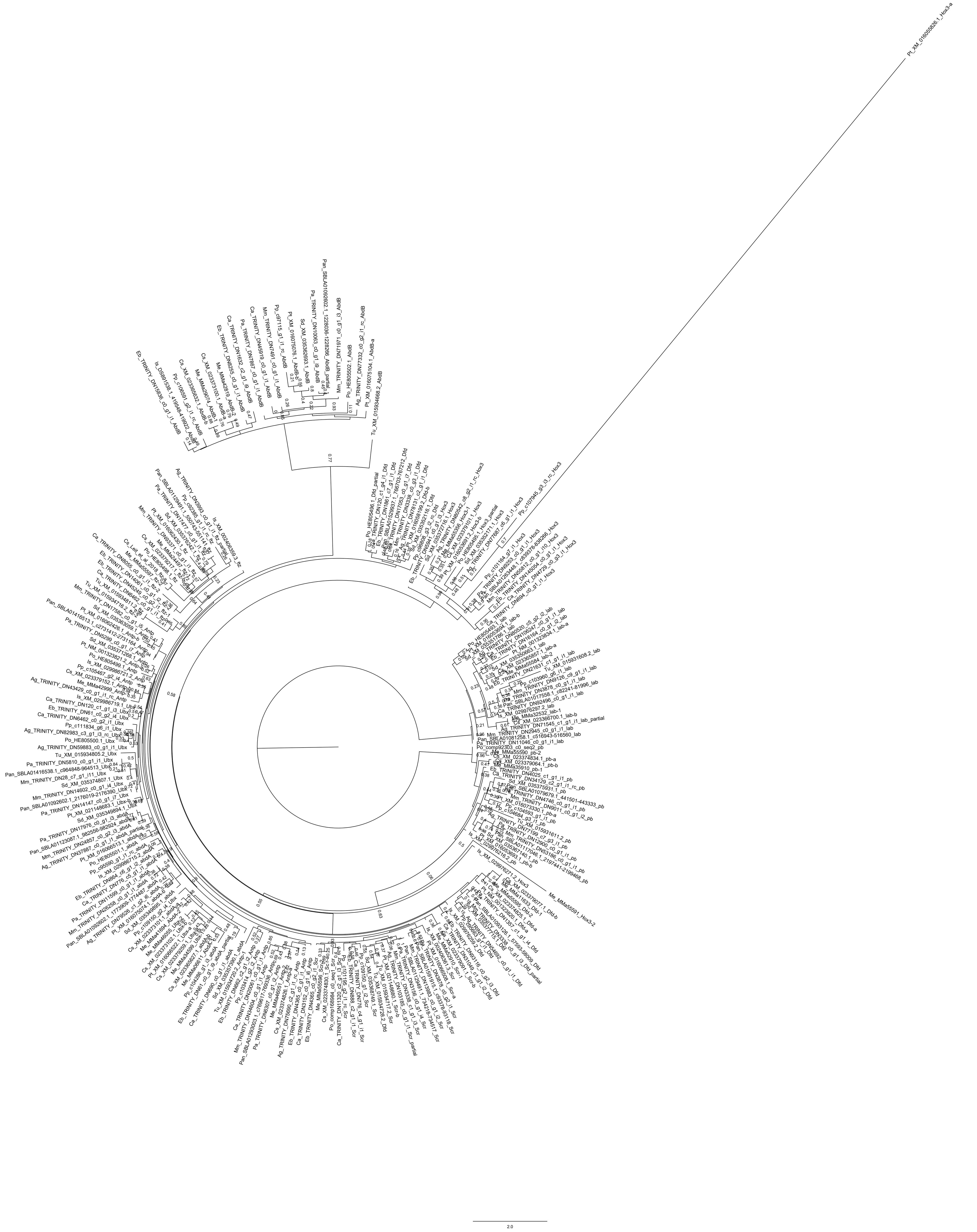
